# Impact of the introgression of resistance loci on agro-œnological traits in grapevine interspecific hybrids

**DOI:** 10.1101/2025.03.31.646340

**Authors:** E. Chedid, C. Rustenholz, K. Avia, V. Dumas, D. Merdinoglu, É. Duchêne

## Abstract

- A major aim in modern grapevine (*Vitis vinifera* L.) breeding programs is the introgression of disease resistance genes along with desired cultural and œnological traits. Understanding the genetic links between resistance genes and agro-œnological traits is a crucial issue for grapevine breeders.
- We studied the genetic determinism of a wide range of agro-oenological traits in a complex interspecific hybrid population and identified two cases of colocalizations with disease resistance quantitative trait loci (QTL).
- The species of origin of chromosomal regions in the off-springs were determined thanks to *in silico* chromosome painting. The linkage drag around resistance genes from primary gene pool was assessed as low due to a good recombination rate with *V. vinifera*.
- We established that wild *Vitis* species are, apart from their interest in disease resistance, essential resources for improving the cultural characteristics of grapevine.

## 1. Introduction

Plant wild relatives have been used in plant breeding programs for decades. They provide the breeders with a wide pool of genetic resources for many traits of interest, in particular tolerance or resistance to abiotic and biotic stresses (Hajjar & Hodgkin, 2007; Prohens et al., 2017). The use of wild species in breeding programs to meet these challenges is howveer often complex, mainly due to the introgression of undesirable traits along with targeted genes. Several studies on annual and perennial plants showed the effect of the introgression of resistance genes from wild species on plant performance (Uauy et al., 2005; Lewis et al., 2007; Danan et al., 2011; Verlaan et al., 2013; Lin et al., 2014; Zhu et al., 2018). For instance, Rubio et al. (2016) found that *Ty-1,* a Tomato Yellow Leaf Curl Virus (TYLCV) resistance gene, decreases tomato yield by 50% when introgressed into traditional Spanish tomato cultivars (*Solanum lycopersicum* L.) and also negatively affects fruit quality. Similar cases were also reported on perennial plants. *Prunus davidiana*, a wild relative of the cultivated peach, is used as a natural resistance source for pest and disease control. Alleles from *P. davidiana* were proved to increase fruit sugar content and red coloration. But several cases of undesirable colocalizations between QTL of resistance to powdery mildew and QTL of agronomical traits, such as fruit and stone size, were identified (Quilot et al., 2004). Breeders generally aim to minimize the percentage of genomic DNA from wild species in an attempt to reduce the risk of linkage drag whether it has positive or negative effect.

In grapevine (*Vitis vinifera* L.), breeding for disease resistance was initiated in Europe after the successive introductions of powdery mildew, phylloxera and downy mildew from North America in the second part of the nineteenth century. The main objective was to combine resistance against phylloxera, powdery and downy mildews as well as producing grapes suitable for wine making. The interspecific hybrids, known as direct producers hybrids, were the first attempt in this direction. They derived from crosses between cultivated European grapevines and wild American *Vitis* species, but they often failed to combine the pleasant flavor of the *V. vinifera* grapes and the resistance of wild species. The poor reputation of these cultivars, some of which carry the undesirable flavors from wild species, undermined further attempts for interspecific hybridization with non-*vinifera* species. However, these hybrids were a starting point for modern breeding programs.

Modern viticulture is facing many challenges, from decreasing phytosanitary treatments to adapting to climate change while preserving wine quality and maintaining production potential. Breeding new grapevine varieties that combine all the desired traits is a powerful lever to meet these challenges which, as in the past, often calls for the introgression of resistance genes originating from wild *Vitis* species into cultivated grapevine (*Vitis vinifera* L.) backgrounds. To develop these new varieties resistant to biotic and abiotic stresses and better adapted to climate change through an optimized selection process, understanding the effect of introgression of resistance genes from wild *Vitis* on agronomical traits is crucial. A recent grapevine study aimed at identifying and quantifying haplotypic blocks of wild ancestry in introgression lines carrying disease resistance QTL (Foria et al., 2022). Nevertheless the effect of linkage drag on grapevine agronomic and oenological aptitudes is still unknown.

The American and Asian *Vitis* species have supplied major resistances genes for fungal diseases such as *Rpv1* (Merdinoglu et al., 2003), *Rpv3* (Bellin et al., 2009; Welter et al., 2007) and *Rpv10* (Schwander et al., 2012) for resistance to downy mildew (*Plasmopara viticola)*, *Run1* (Barker et al., 2005) and *Ren3/9* (Welter et al., 2007; Zendler et al., 2017) for resistance to powdery mildew (*Erysiphe necator*), and *Rgb1* (Rex et al., 2014) for resistance to black rot (*Guignardia bidwellii*).

Recent studies in grapevine aimed at identifying and quantifying the haplotypic blocks of wild ancestry in introgression lines carrying disease resistance QTL. Foria et al. (2022) found that newly released resistant varieties contain 76.5–94.8% of *V. vinifera* DNA and that the linkage drag of wild alleles around known resistance genes varies between 7.1–11.5 Mb. But the effect of this linkage drags on grapevine agronomical and oenological aptitudes is still unknown, thus the traits that can be inherited from wild species while introgressing resistance genes is a crucial question to address.

The present study aims at identifying colocations of resistance genes with other traits of interest and determining their origins in an attempt to avoid undesirable linkage in future crosses. A wide range of phenotypic traits was analyzed in the vineyard thanks to a progeny from a cross between two complex grapevine interspecific hybrids, involving different origin of *Vitis* species and carrying each several factors of resistance to fungal diseases. QTL detection of agro-oenological traits was based on high-density genetic maps. Using an *in silico* chromosome painting approach, the species of origin of haplotypic blocks in the progeny were identified. All together, these results allowed us to provide new insights into the effects of the introgression of resistance genes on agro-oenological traits.

## 2. Materials and methods

### 2.1. Plant material and experimental design

We have chosen to study a progeny, called ‘50025’, obtained as part of the French INRAE-ResDur grapevine breeding program that aims at creating varieties combining resistances to downy and powdery mildew with berry composition suited to the production of high quality wines. Population ‘50025’ is the result of a cross between two interspecific hybrids, IJ119 and Divona, each with distinct ancestors and resistance gene combinations. The 50025 pedigree is complex and includes, in addition to cultivated grapevine (*V. vinifera)*, *V. aestivalis*, *V. berlandieri*, *V. cinerea*, *V. labrusca*, *V. lincecumii*, *V. riparia*, *V. rupestris, V. rotundifolia* and *V. amurensis* (Figure 1, Figure S1). The 249 individuals of population ‘50025’ display various combinations from 0 to 7 resistance loci. A set of 209 genotypes out of 249 were grafted on SO4 rootstock in 2015 and planted in 2016 in the INRAE vineyard in Colmar, France (48.1 ° N, 7.33 °E) at a density of 4200 plants per ha according to a randomized complete 2-block design, including the parents, and the control variety *V. vinifera* cv. Chardonnay. Each genotype was represented by one elementary plot of three plants per block. The control variety, Chardonnay, was represented by six elementary plots per block.

**Figure 1.**
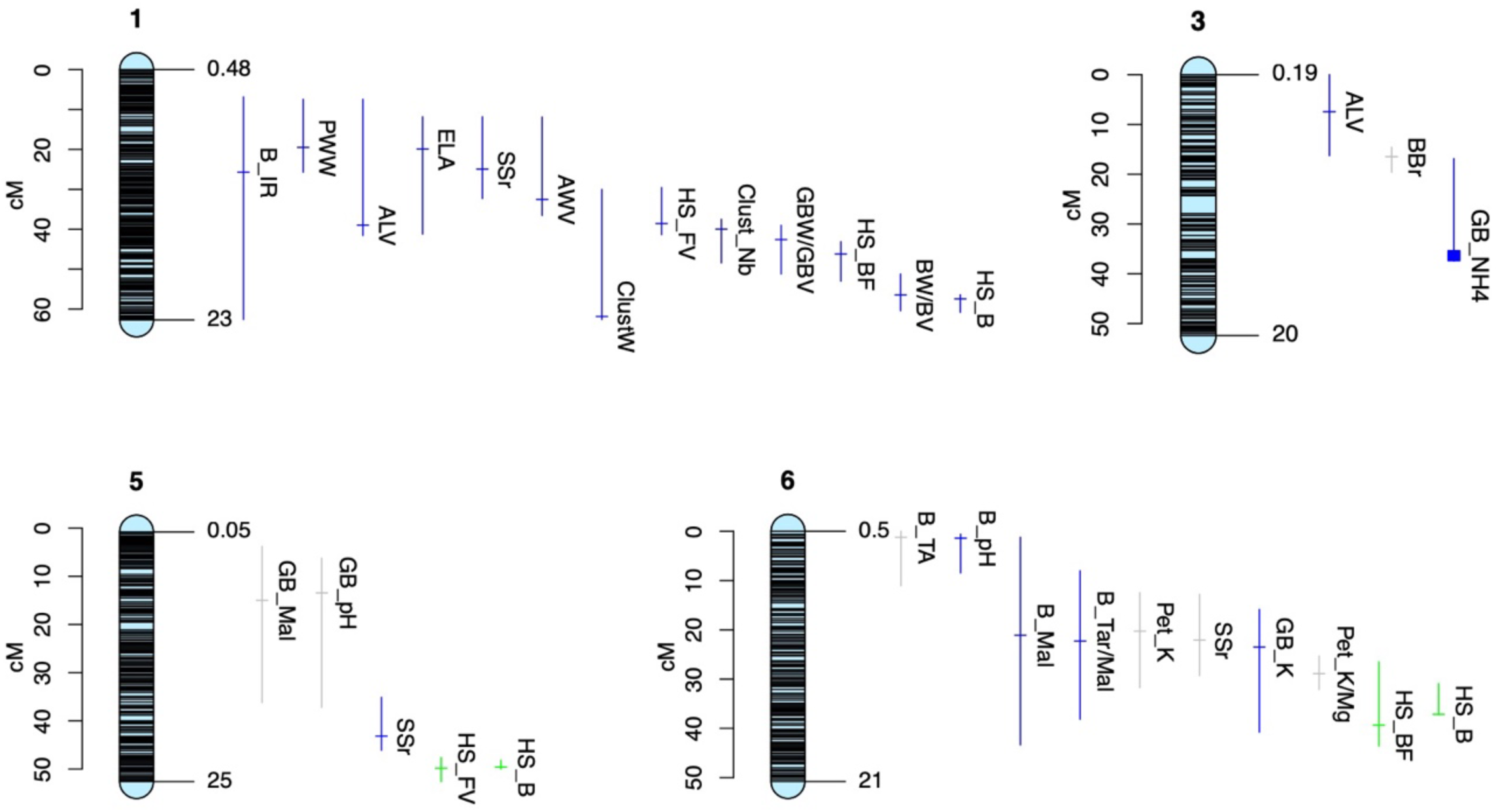
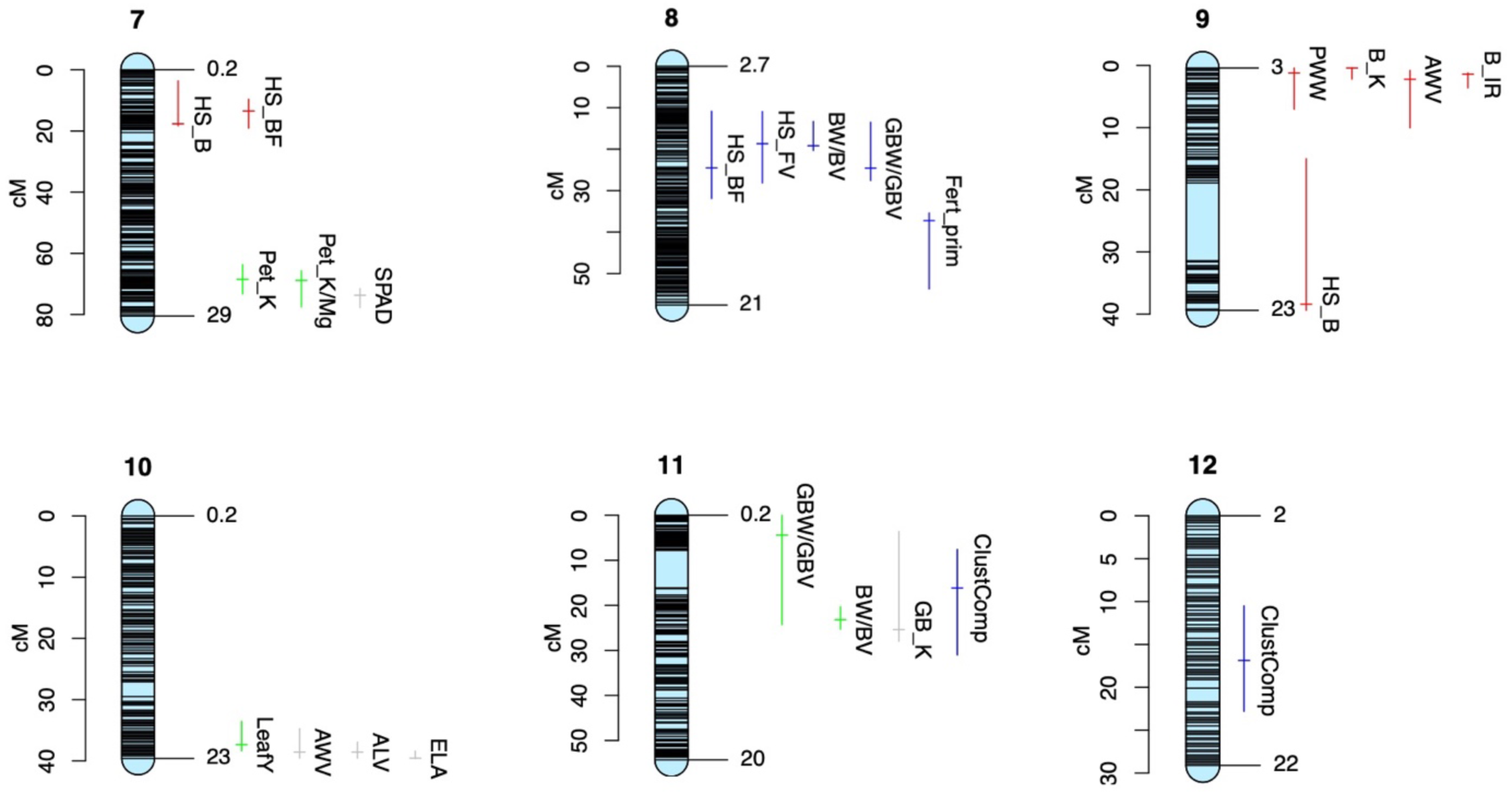
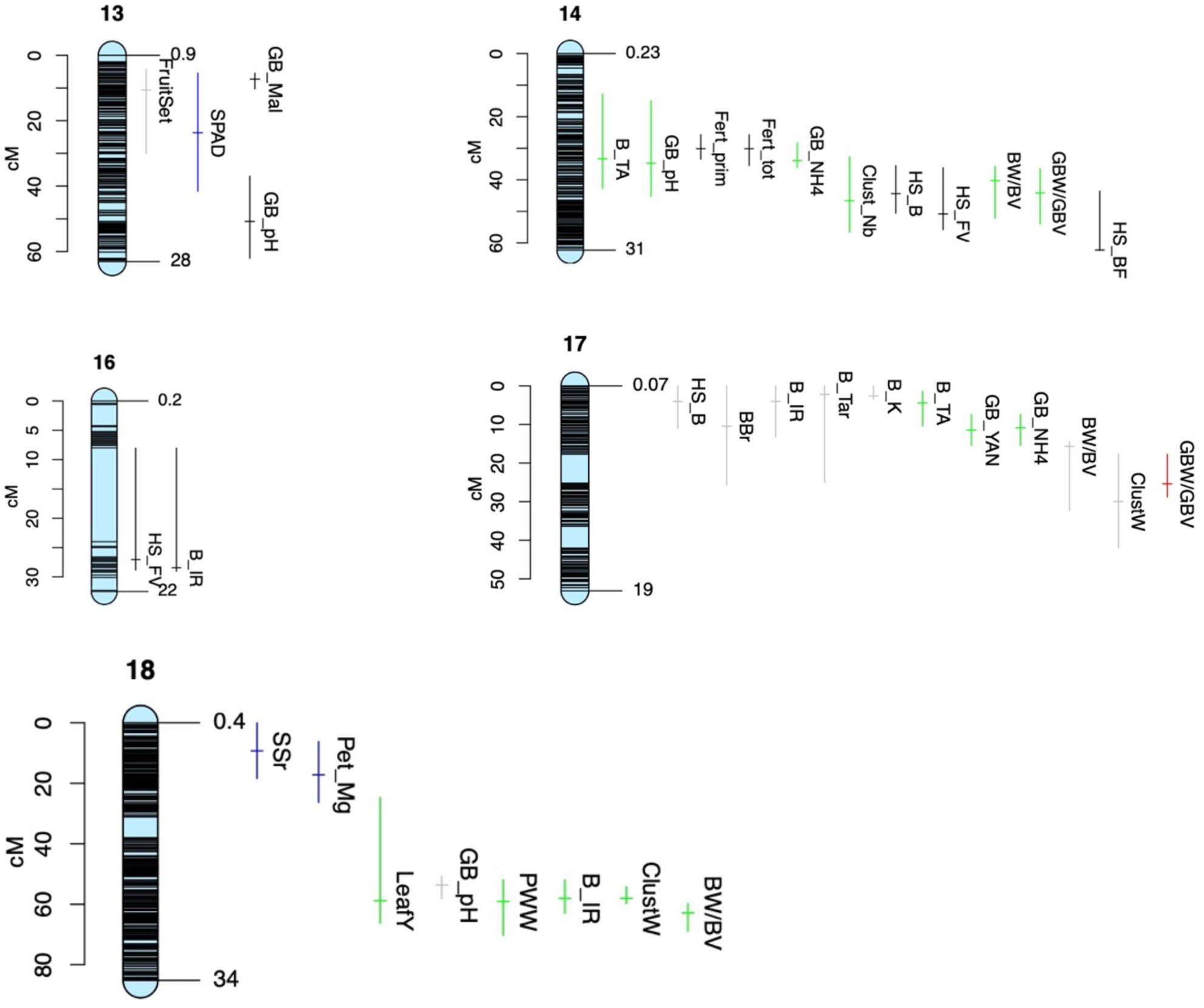
QTLs of agronomical traits detected on the consensus map of the population. Based on ‘chromosome painting analysis’, blue indicates *V. vinifera* origin, green indicates American *Vitis* origin, red indicates Asian *Vitis* origin, grey indicates several species origin and black indicates unidentified origin.

### 2.2. Genotyping and genetic maps

Genotyping-By-Sequencing (GBS) technology was used for genotyping the population ‘50025’ and both parents (Elshire et al., 2011). GBS banks were sequenced on an Illumina HiSeq4000 platform (paired-end, 2x 100 bp). The reads were aligned on the *V. vinifera* PN40024.v4 (Velt, 2022) reference genome using `BWA`. SNP calling was performed using the `gstacks` command of Stacks v2 pipeline (Catchen et al., 2013). The output file in Joinmap format was filtered to only keep the most informative and reliable markers. Briefly, SNPs with more than 10% missing data, with non-Mendelian segregation (χ2 test, P < 0.05) or not consistent with the genotype of the parents, were discarded. Since the progeny in study is a breeding population, genotyping data of eleven microsatellite markers were used in Marker Assisted Selection (MAS) to identify the genotypes carrying the resistance loci. The genotyping data of the SSR markers were added to SNP data from GBS and used for the construction of genetic maps. Genetic maps were constructed using Lep-Map3 (Rastas, 2017). `ParentCall2` module of Lep-Map3 was used to call parental genotypes, `SeparateChromosomes2` module was used to split the markers into 19 linkage groups (LGs) and `OrderMarkers2` module was used to order the markers within each linkage group using 30 iterations per group and finally computing genetic distances. The output data were converted into R/qtl format for R software (Broman et al., 2003). The consensus map was coded as a 4-way cross for R/qtl with 4 parental alleles : A, B from IJ119 and C, D from Divona.

In order to identify the parental origin of chromosomal regions in the progenies and thus identify the species of origin, it is mandatory to have the phasing of each marker to assign each nucleotide to the corresponding haplotype of the parent. While this information was not available from Lep-Map3 genetic maps, we generated parental genetic maps using `ASMap` function of R software. The output vcf file of `gstacks` was filtered to keep reliable and informative markers. To keep the highest number of markers without affecting maps quality, we kept markers with up to 30% missing data. The script used for map construction includes a coding system where each SNP was coded in the two possible phases. Out of the 38 LGs calculated through each parental map construction, we chose 19 of them. The phased output data from parental maps, with the reference and alternative nucleotide for each SNP, were used later for “chromosome painting analysis”.

### 2.3. Phenotyping

Forty-one traits related to phenology, yield, vegetative development, and berry quality were measured for three consecutive seasons (2019 to 2021) on the IJ119 x Divona progeny in the field. All variables and their abbreviations are presented in table S1.

Three phenological periods were studied according to heat sum system described in Duchêne et al. (2010). We calculated for each genotype, the heat sum between 15 February and budburst, between budburst and flowering and between flowering and véraison.

The traits of vegetative development - exposed leaf area, pruning wood weight, apparent canopy volume, apparent wood volume and chlorophyll content were obtained according to Chedid et al. (2022-submitted).

Potassium and magnesium content in petioles were measured on 24 petioles per genotype. The sampling was conducted around véraison time: petioles were sampled, dried then grinded and used for potassium and magnesium quantification using ICP-AES method.

For each plant, we counted the number of primary and secondary shoots and their respective number of inflorescences. Primary fertility is the ratio between the number of primary inflorescences and the number of primary shoots. Total fertility is the ratio of total inflorescences on primary shoots. Then, at harvest, we counted the number of clusters harvested and measured their weight.

Fruit set quality was evaluated per plant on a visual scale from 0 (poor) to 3 (good). Assessment of bunch compactness was done around maturity according to the OIV 204 scale, from 1 (very loose) to 9 (very compact) (OIV, 1983).

Samples of 200-mL box of berries from both parents, progenies of Divona x IJ119, and Chardonnay variety were obtained at two dates. The first sampling was performed independently for each genotype during the véraison process, and only green and hard berries were picked. The second sampling was as well performed independently for each genotype, when it accumulated 380 degree.days (dd) (Tmean, base 10) after véraison (50% of soft berries). For both types of samples, the berries were counted and weighed.

Green berries were ground with an electric blender. The pH of berry juice was measured with a 340i pH-meter (WTW, Weilheim, Germany) calibrated daily.

Berry samples at 380 dd were ground with an automatic sieve (Robot Coupe C80, Montceau-en-Bourgogne, France). Titratable acidity, pH, and total soluble solids (TSS, °Brix) were measured on the juice. Total acidity was measured on 5 mL of clear juice using the titrator (Titroline Easy) with NaOH (0.1 N). TSS was measured with a PAL-1 refractometer (Atago®,PAL-1).

Green berry juices and berry at 380 dd were diluted in distilled water with a coefficient of 1 to 10 and 1 to 4, respectively. Samples were frozen in order to avoid precipitation of organic acids and subsequently analyzed.

The concentration of malic acid in berries was determined with an enzymatic method and the concentration of tartaric acid with a colorimetric method using ammonium metavanadate (Microdom, Taverny, France). Both acids were analyzed in duplicate on a Lisa 300 automatic analyzer (Biocode Hycel, Le Rheu, France).

The concentration of potassium in berries was determined with flame spectrophotometry.

The concentration of ammoniacal nitrogen in green berries was determined with an enzymatic method and the concentration of alpha-amino nitrogen was determined with a colorimetric method using o-phtaldialdehyde (Oenolab). Yeast assimilable nitrogen in green berries is the sum of ammoniacal and alpha-amino nitrogen content.

### 2.4. Statistical analysis

For each trait, we inspected the raw phenotypic data for non-normal distribution, using a Shapiro-Wilk normality test. The best transformation, according to `bestnormalize` function of R, was computed for datasets not satisfying normality criteria. Then, we fitted the following linear mixed model: *y_ij_ = μ+ G_i_ + Y_j_+* ε*_ij_* where *μ* is the mean, *G* and *Y* are the random effects of the genotype and the year, respectively. Best Linear Unbiased Predictors (BLUPs) values were extracted from anovas during the process (`anova` function in R).

Calculating the heritabilities with the conventional method using the genetic and environmental variances from the anovas on the segregating population was not possible because of the absence of biological replicates. We estimated the environmental variance α_e_^2^ using the six plots of Chardonnay planted across the trial. The genetic variance was calculated by subtracting α_e_^2^ from α_t_^2,^ the variance observed over all the genotypes from the 50025 progeny. Then broad-sense heritabilities were calculated as H^2^ = (α_t_^2^ -α_e_^2^)/ α_t_^2^.

### 2.5. Interval mapping analysis

Quantitative trait loci (QTL) detection was performed on the consensus map with the R/qtl software (Broman et al., 2003) using the multiple imputation method (“draws” = 64) and the one-dimension scan command `scanone`. LOD (logarithm of odds, that evaluates the likelihood of the presence of a QTL) significances were ensured with permutation tests (1000 permutations). QTL models were constructed step-by-step after the refinement of the QTL position (`refineqtl`), and the search for supplementary QTLs (`addqtl`). The LOD score and the percentage of variance explained by a QTL in a QTL model were assessed with analyses of variance using type III sums of squares (`fitqtl`). Confidence intervals were calculated as Bayesian credible intervals (`bayesesint`) with a probability of coverage of 0.95 Finally, all pairwise interactions possible in the multiple QTL model were added when significant with the`addint` function.

QTL detection was performed on annual data and BLUPs of three years calculated from linear mixed models. To simplify the results, we show data for QTL models for the BLUPs that include only QTL that where considered as stable over years. A QTL was considered stable if it was significant on at least two years and on BLUPs.

### 2.6. Chromosome painting

“Chromosome painting analysis” consists of identifying the origin, in terms of ancestry or geographical source, of chromosomal regions in the genome of an individual (Figure S2). Thanks to genetic maps, which allow the phasing of the SNPs, and the genome sequencing from parents, grandparents and/or accessions from various geographical sources, the parental or geographical origin of each allele in the offspring can be identified. By examining the adjacent SNPs in blocks, the origin of chromosomal regions can be determined.

The SNPs and alleles of the individual of interest were compared with a genotyping file, i.e. a vcf file obtained through SNP calling analysis using resequencing data. In our study, 135 vine accessions representative of several *Vitis* species were used and grouped according to their species and their geographical origin: *Flexuosae* (10 accessions), *Spinosae* (2 accessions) and *V. coignetiae* (3 accessions) for Asian *Vitis*, *Ripariae* (18 accessions), *Cinereae* (13 accessions), *V. labrusca* (6 accessions), *Candicansae* (3 accessions), *V. cordifolia* (4 accessions), *V. arizonica* (1 accession) and *Aestivalae* (8 accessions) for American *Vitis*, *V. vinifera* (54 accessions) for European *Vitis* and *V. rotundifolia* (7 accessions) for the *Muscadinia* group. For each allele, its rate of presence was calculated in each species/geographic group. A segmentation analysis was then run on each haplotype to identify blocks of consistent origin. The species / geographical group with the greatest average presence rate on the block was chosen as the origin of the corresponding chromosomal region. Finally, both “painted” karyotypes could be obtained for the individuals of interest. If a segregating population is to be painted, the chromosome painting strategy is first applied on the parents of the cross and the origins are then transferred to the progenies according to the recombination breakpoints. In this way, the origin of the two haplotypes of each chromosome is identified for each offspring. Each haplotype is divided into haplotypic blocks according to the species of origin. Homologous regions are the chromosomal regions with both haplotypes from the same origin while homeologous regions are sequences with haplotypes from different origin.

## 3. Results

### 3.1. Heritabilities and segregation

The statistics for all the variables, for Chardonnay, the parents and the progeny are presented in table S2. Overall, the studied traits showed good level of diversity in the population and several variables showed transgressive segregations.

Heritabilities calculated for all studied traits per year are presented in table S1. For phenological periods, heritabilities were high especially for period between flowering and véraison time (>0.8). For growth traits, important variations between different seasons were observed for all variables except the LiDAR data. For yield components, primary fertility (0.96 in 2020) and green berry weight (0.92 in 2021) had the highest heritabilities. The heritability of green berry pH was the most stable and the highest among berry quality traits (0.83 in 2020 and 0.97 in 2021). The concentration of malic acid in berries had the lowest heritability in 2020 (0.15). To summarize, we observed for most of the studied traits a wide variation that is due to genetic effect. In addition, the transgressive phenotypes observed for certain variables indicate that both parents are involved in the genetic variability of these traits, *i.e.* they may both be carriers of favorable or unfavorable QTLs. These results allowed us to proceed with genetic and QTL analyses.

### 3.2. Genetic maps

Genetic information from 249 genotypes of the population was used for the construction of genetic maps. Coverage rate is the ratio of physical distance between distal mapped markers and the total physical chromosome length. The average coverage rate of genotyping data on all chromosomes is 99.98%. The three maps, constructed with Lep-Map3, have 19 LGs and a high marker density of 0.1 cM. The consensus map, with a total length of 1017.3 cM, contains 28 207 SNP. The female map has 19 930 SNP with a total genetic length of 1105.3 cM and the male map has 20 093 SNP covering 1100 cM. Through the construction of genetic maps, the coverage rate decrease due to filtering of non-informative SNP (homozygous, distorted…). IJ119 covers 88.5% (min 44.6% on LG5-max 99.43% on LG18) of the total genome and Divona covers 85.5% of the genome (min 11% on LG2, max 99.4% on LG8). The SSR markers associated with QTL of resistance are used for MAS as a criterion for the selection of resistant genotypes. We integrated the SSR genotyping data to identify the regions carrying the resistance loci. Positions of SSR markers on genetic maps are presented in table 1. Genetic and physical distances of markers in the maps were highly correlated (>0.9) (Figure S3) which is essential for the detection of QTL.

**Table 1.**
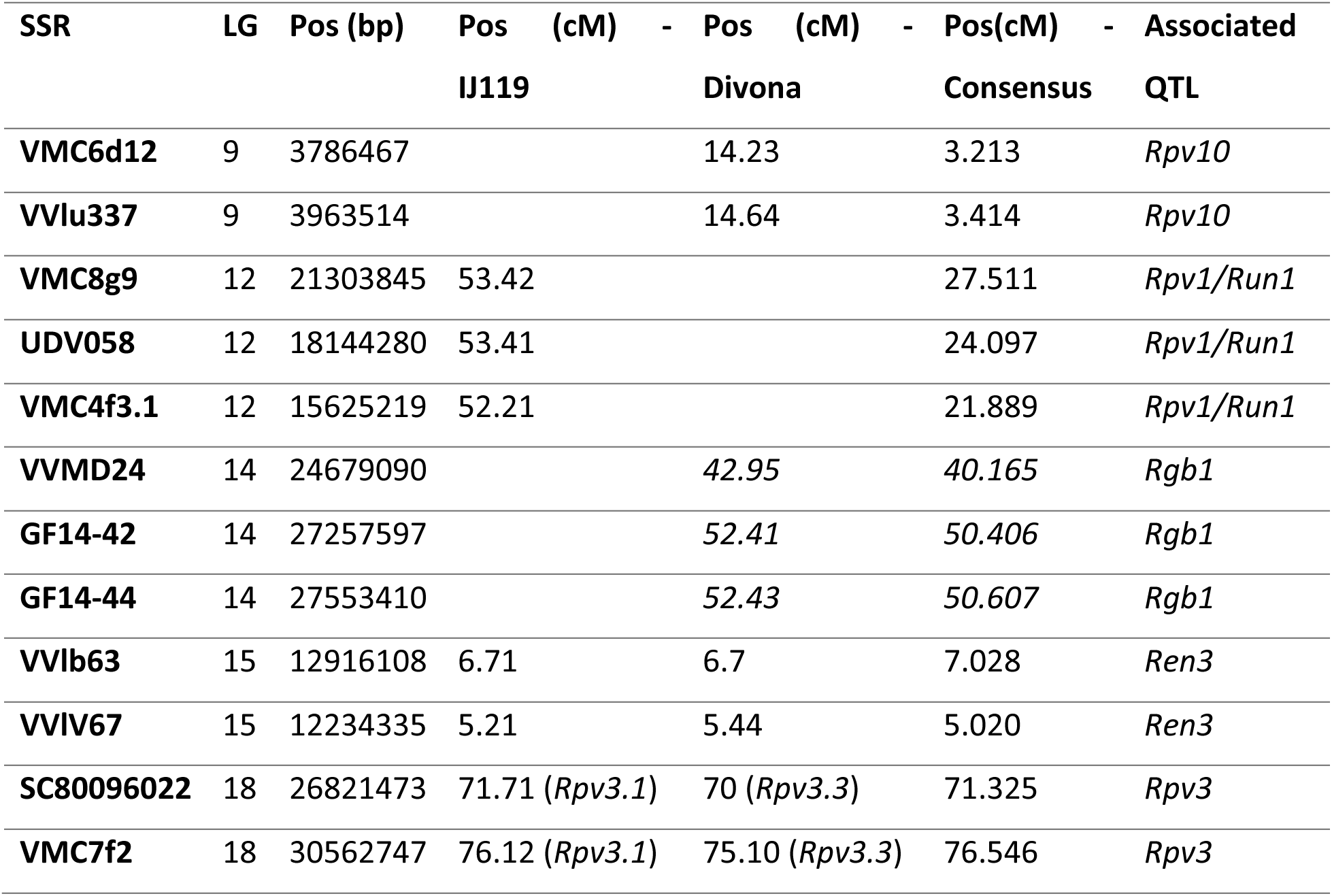
Positions of SSR markers associated to resistance genes on the genetic maps of the IJ119xDivona population.

| SSR | LG | Pos (bp) | Pos (cM) | - | Pos (cM) | - | Pos(cM) | - | Associated |
| --- | --- | --- | --- | --- | --- | --- | --- | --- | --- |
|  |  |  | IJ119 |  | Divona |  | Consensus |  | QTL |
| **VMC6d12** | 9 | 3786467 |  |  | 14.23 |  | 3.213 |  | *Rpv10* |
| **VVlu337** | 9 | 3963514 |  |  | 14.64 |  | 3.414 |  | *Rpv10* |
| **VMC8g9** | 12 | 21303845 | 53.42 |  |  |  | 27.511 |  | *Rpv1/Run1* |
| **UDV058** | 12 | 18144280 | 53.41 |  |  |  | 24.097 |  | *Rpv1/Run1* |
| **VMC4f3.1** | 12 | 15625219 | 52.21 |  |  |  | 21.889 |  | *Rpv1/Run1* |
| **VVMD24** | 14 | 24679090 |  |  | 42.95 |  | 40.165 |  | *Rgb1* |
| **GF14-42** | 14 | 27257597 |  |  | 52.41 |  | 50.406 |  | *Rgb1* |
| **GF14-44** | 14 | 27553410 |  |  | 52.43 |  | 50.607 |  | *Rgb1* |
| **VVlb63** | 15 | 12916108 | 6.71 |  | 6.7 |  | 7.028 |  | *Ren3* |
| **VVIV67** | 15 | 12234335 | 5.21 |  | 5.44 |  | 5.020 |  | *Ren3* |
| **SC80096022** | 18 | 26821473 | 71.71 (*Rpv3.1*) |  | 70 (*Rpv3.3*) |  | 71.325 |  | *Rpv3* |
| **VMC7f2** | 18 | 30562747 | 76.12 (*Rpv3.1*) |  | 75.10 (*Rpv3.3*) |  | 76.546 |  | *Rpv3* |

The phase of each SNP in the genetic maps is an essential information to be used for chromosome painting analysis. So we constructed genetic maps using `ASMap` in order to have the phasing of all loci. Both parental genetic maps constructed with `ASMap` were less dense than genetic maps of Lep-Map3 since markers with the same pattern of heterozygosity in both parents (ex: ABxAB) are not supported by `ASMap`. IJ119 map have 12002 markers covering 4187.6 cM with 0.3 cM marker density. Divona map had 11048 markers covering 4219.6 cM and 0.4 cM density. Genetic and physical distances of markers are consistent with those obtained with Lep-Map3 maps although some chromosomal regions like LG15 were less covered due to high percentage of hkxhk markers (data not shown).

### 3.3. QTLs of agronomical traits

A wide phenotyping program was conducted for three consecutive years and covered most of the agronomical traits that can be interesting in grapevine breeding programs. More than 90 QTLs located on 16 LGs were detected for 34 variables. Four berry composition-related traits had no QTL identified: GB_Tar, B_NH4, B_Ca and B_Mg. Four LGs (LG1, LG6, LG14 and LG17) carry each more than ten QTLs, accounting for half of the total. On the contrary, some others (LG3, LG12 and LG16) bear only very few or even none (LG2, LG4 and LG19). All QTL detected on the consensus map are presented in Figure 1 and Table S1.

Thirteen QTLs were detected on LG1 for phenological stages, yield component, plant vigor, and berry quality. These QTLs are divided into two clusters. QTLs of phenology, berry weight, and number of clusters per plant are linked together. The allelic combination that increasesand HS_FV, increases BW and decreases the Cluster_Nb. The second cluster of QTLs highlights the link between plant vigor and sugar content in berries. QTLs of ELA and PWW overlap with QTL of TSS of the berries. The allelic combination that increases leaf area also increases berry sugar content. The colocalization between these QTLs is probably due to the physiological link between them as a larger canopy increases the photosynthetic capacity of the plant and potentially sugar accumulation in the berries.

Ten QTLs were detected on LG6 for phenology, mineral nutrition, and berry acidity. Confidence intervals of GB_K, B_Tar/Mal, B_Mal, Pet_K/Mg and Pet_K overlap, suggesting a link between these traits. As shown in Figure S4, the same allelic combination increases GB_K, B_Tar/Mal, and pH, and decreases B_Mal and B_TA. Potassium in berries partially neutralises organic acids, which reduces berry acidity.

On LG14, the allele D of Divona increases number of clusters per plant and decreases GB_W and BW. But the maternal allele A increases BW. Three QTLs of B_TA, GB_pH and GB_NH4, coming from both parents, colocalize on LG14. The maternal allele A increases GB_NH4, GB_pH and decreases B_AT. The paternal allele C increases GB_NH4 but increases B_TA and GB_pH (Figure S5).

On LG17, we detected QTLs for yield components and berry composition. QTL of GBW comes from Divona while QTL of BW and ClustW comes from both parents: the paternal allele C increases GBW, BW and ClustW, the maternal allele A increase only BW and ClustW. CI of BW and ClustW overlap with QTL of GB_NH4 and GB_YAN (Figure S6). The maternal allele A increase nitrogen content in berries which may explain the effect of maternal allele on berry weight. On the other side, QTL of nitrogen content overlap with QTL of B_TA. QTLs of berry acidity comes from both parents, the allelic combination AD decreases B_AT, B_Tar and B_K.

The colocalization between QTLs of different variables suggests that these traits may be linked in three different possible ways: i) different genes with close physical position on the genome, ii) a pleiotropic effect of the same gene or ii) a physiological link between different traits leading to the colocalization of QTLs. Berry weight and phenological stages appear to be linked on different LGs. QTLs of berry weight are linked to QTLs of budburst on LG1 and 14. Also, on LG8 the QTL for heat requirements between budburst and flowering is linked to berry weight QTL. In all three cases, earlier phenological development is linked to smaller berries. The fact that it has already been observed that the late ripening cultivars have larger berries compared to early ripening cultivars (Clingeleffer & Davis, 2022) supports the hypothesis of a physiological link.

We noticed a link between the QTL for secondary shoots ratio and QTLs of phenological stages. On LG 1, 5, and 6 there are QTLs for both SSr and phenological stages and in all three cases, the genotypes with earlier phenological development have a higher secondary shoots ratio. Also, the QTLs of SSr on LG6 and 18 are linked to QTL of mineral nutrition, potassium and magnesium in petioles. Lower SSr in plant results in higher concentration of K or Mg in petioles.

Four QTLs of potassium content in berries were detected on LG 6, 9, 11 and 17 but only QTL on LG6 is apparently linked to potassium content in petioles.

On LG7, QTL of SPAD and Pet_K colocalize. SPAD is correlated to photosynthetic capacity of the leaves. Petrie et al. (2000) investigated grapevine photosynthesis and found some evidence of the potential relationship between leaf photosynthetic activity and K export from/import to leaves. In addition, potassium deficiency affects the photosynthetic activity due to dramatic decrease in leaves chlorophyll content (Zhao et al., 2001).

The QTLs of nitrogen content in berries on LG14 and 17 colocalize with QTLs of acidity. This result support also the association between organic acids in berries and yeast assimilable nitrogen found by Clingeleffer & Davis (2022). According to the allelic effect of these loci, nitrogen and acidity in berries vary in opposite directions. Clingeleffer and Davis (2022) reported significant negative correlation between pH and YAN in berries which indicates that soluble amino acids can have a role in berries acid balance. Nitrogen content in green berries seemed to have an effect on berry weight.

### 3.4. Colocalization of disease resistance factors with QTLs of agronomical traits

Seven known disease-resistance genes segregate in the studied population on LG9, 12, 14, 15, and 18. We studied the genetic links between these loci and other QTL of agronomical traits on the same LGs. In order to be more accurate, we compared positions and allelic effects on the parental maps instead of the consensus map (Figure 2).

**Figure 2.**
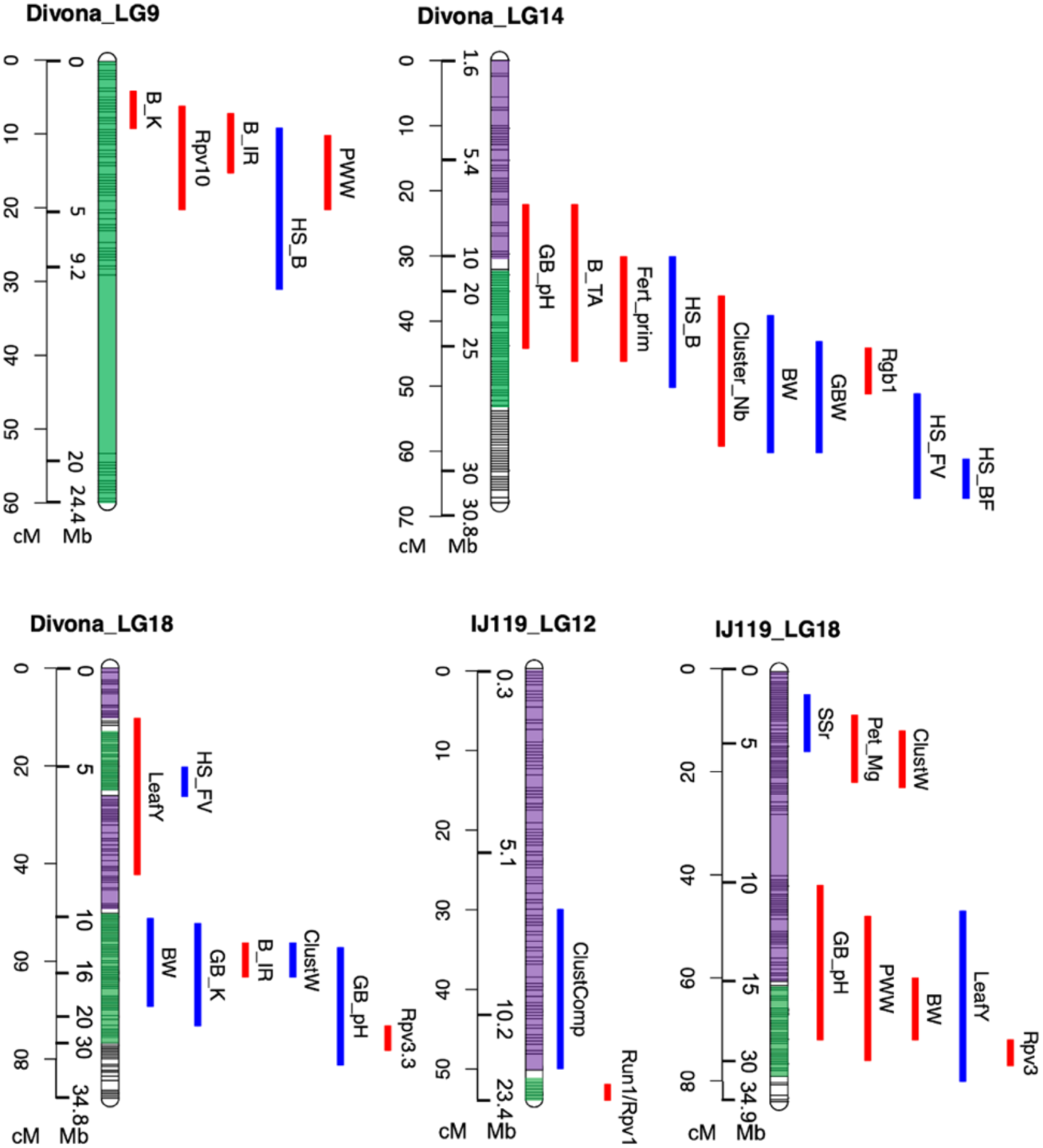
QTLs positions of agro-oenological traits on LGs carrying resistance genes. Green color on LGs = homeologous regions. Purple on LGs = homologous regions (*V. vinifera*). Red QTL = trait value increases with resistance. Blue QTL = trait value decreases with resistance.

VMC6d12 and VVlu337 are associated to *Rpv10*, they are known to be downstream of the resistance loci on LG9. Accordingly, *Rpv10*, QTLs for pruning wood weight, and TSS in berries colocalize. Thus, the same haplotype conferring resistance to downy mildew increases the pruning weight and TSS in berries. Two other QTLs, for potassium content in berries (1490805 bp) and budburst date (4819823 bp), are found in the same region of *Rpv10* and their confidence interval overlap; so there is a high probability of having higher potassium content in the berries and earlier budburst date associated with *Rpv10*. As LG9 of Divona is entirely homologous, *Rpv10* and the linked QTLs are from the same origin.

On LG14 of Divona map, 9 QTLs of yield, berry quality and phenology were detected in the same region as *Rgb1*. The QTLs of berry weight and number of clusters (26.085.884 bp) colocalize with *Rgb1* (24-27Mbp). Those three QTL are in the homeologous region and have the same origin. It is previously reported that earlier phenological development results in smaller berries which is the case on LG14 where three QTL of phenological stages are detected in the same chromosomal region and are obviously linked to berry weight. The QTL of berry acidity and pH are highly linked with *Rgb1.* The resistance to BR here is linked to earlier genotypes, higher fertility, and smaller and more acid berries.

On LG12 of IJ119, only one QTL for bunch compactness was detected next to *Rpv1* and *Run1* but not on the same haplotypic bloc. The QTL of ClustComp and *Rpv1/Run1* have different origins. Resistance to mildews in this population only is associated with lower bunch compactness.

On LG18 of IJ119, four QTL for BW, PWW, LeafY, and GB_pH were detected in the same region of *Rpv3.1* and have a high probability to segregate with the resistance loci in this population. The QTL of LeafY, PWW and GB_pH have large confidence intervals extended on both homologous and homeologous regions of the chromosome and could not be associated to one species.

On the other hand, on LG18 of Divona, *Rpv3.3* and five other QTLs are detected in the same homeologous region. According to allelic effects, there is a high probability of having lower berry pH, yield, berry weight, potassium content, and higher sugar content in berries in the progenies carrying *Rpv3.3*.

According to the SSR markers (VVIb63 & VVIv67) positions on the consensus map, *Ren3* is located between 5 and 7cM. There was no QTL detected in this region

### 3.5. Donor species and linkage drag of disease resistance loci

Haplotypic blocks of seven species (*V. vinifera, V. amurensis, V. coignetiae, V. riparia/V. rupestris, V. labrusca, V. aestivalis, V. rotundifolia*) were identified in the parents and the progeny. The species from which the haplotypic blocks of both parents originate are shown for each chromosome pair in Figure 3. IJ119 display around 90% *V. vinifera* genome: nine chromosome pairs are detected as entirely homologous, ten chromosomes have homeologous regions and the *Ripariae* group is the most present wild (or non-European) *Vitis* species. Only one homeologous region *V. vinifera - V. rotundifolia* is identified on LG12 and one homeologous region with Asian *Vitis* is identified on LG18. Several blocks could not be associated to any species because of low marker density especially on LG15. Divona have less *V. vinifera* genome (80%) compared to IJ119. Nine chromosomes pairs have regions from American species while three from Asian species. Large haplotypic blocks could not be associated to any species on LG2, 15 and 16.

**Figure 3.**
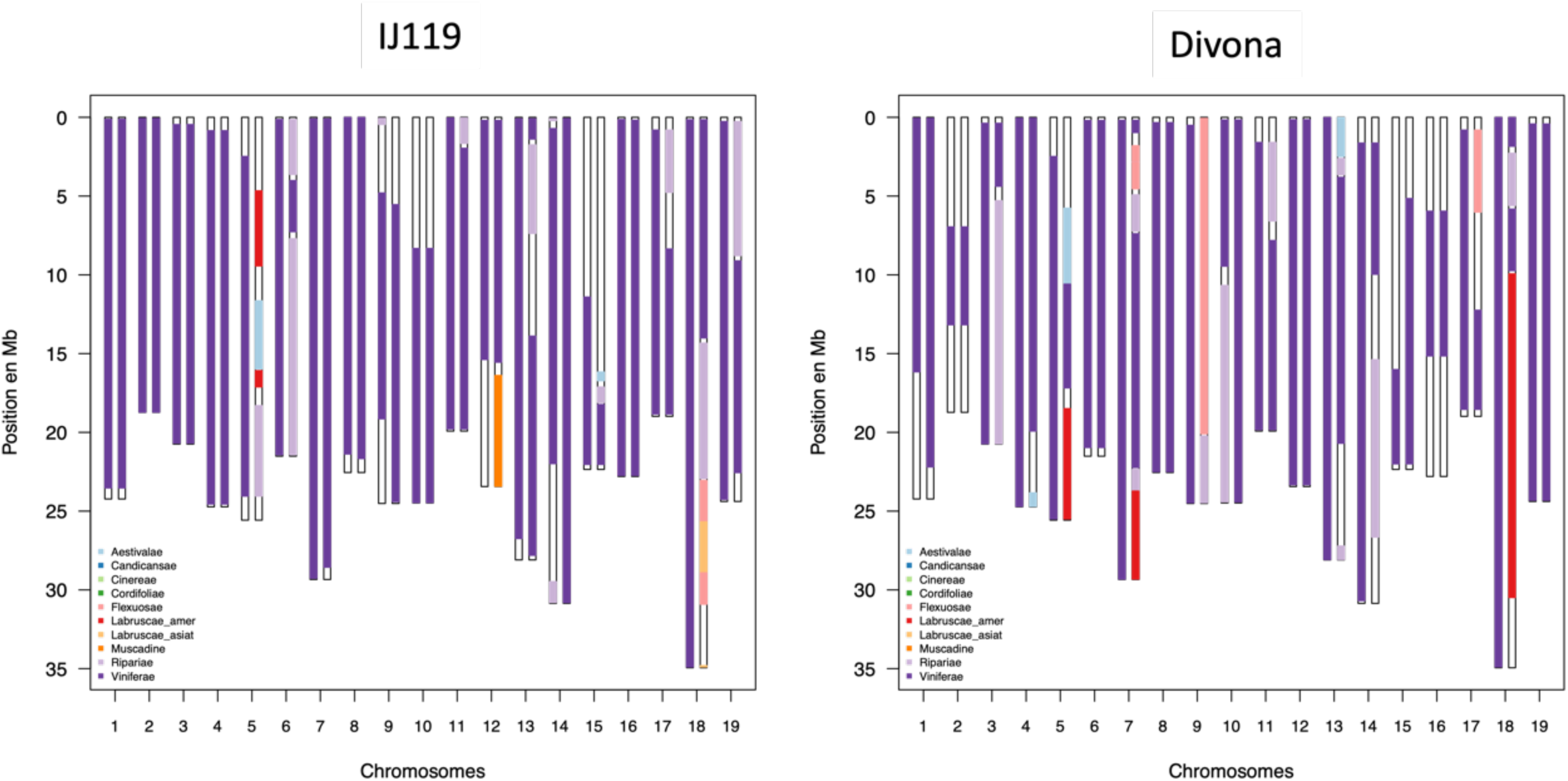
Species of origin of haplotypic blocks on 19 LGs of both parents. Blue = *V. aestivalis* (American), Pink = *V. amurensis* (Asian), Red = *V. labrusca* (American), Orange = *V. rotundifolia* (Muscadinia), Yellow = *V. coignetiae* (Asian), Light Purple = *Ripariae* (*V. riparia* / *V. rupestris*) (American), Purple = *V. vinifera* (European) and White = Undetermined.

We identified the haplotypic blocks carrying the resistance loci introgressed from the wild species, in both parents and offspring, the positions of known resistance factors being determined thanks to the associated SSR markers (Figure 4). Then, we determined, for each resistance locus, the smallest haplotypic block transmitted to the resistant individuals of population ‘50025’. *Rpv1/Run1* locus on LG12 is located in the homeologous block of *V. vinifera - V. rotundifolia* (Figure 4A) confirming that *V. rotundifolia* is the donor species of this resistance locus. This 7.3Mbp haplotypic block recombines very weakly with the homeologous *V. vinifera* bloc: it corresponds to a genetic distance of less than 2 cM, whereas, on the homologous region of the same chromosome, 15 Mbp leads to a distance of 50 cM, i.e. a recombination rate increased by more than a factor of ten. This haplotypic block appears to be preserved from recombination in all genotypes carrying the resistance locus.

**Figure 4.**
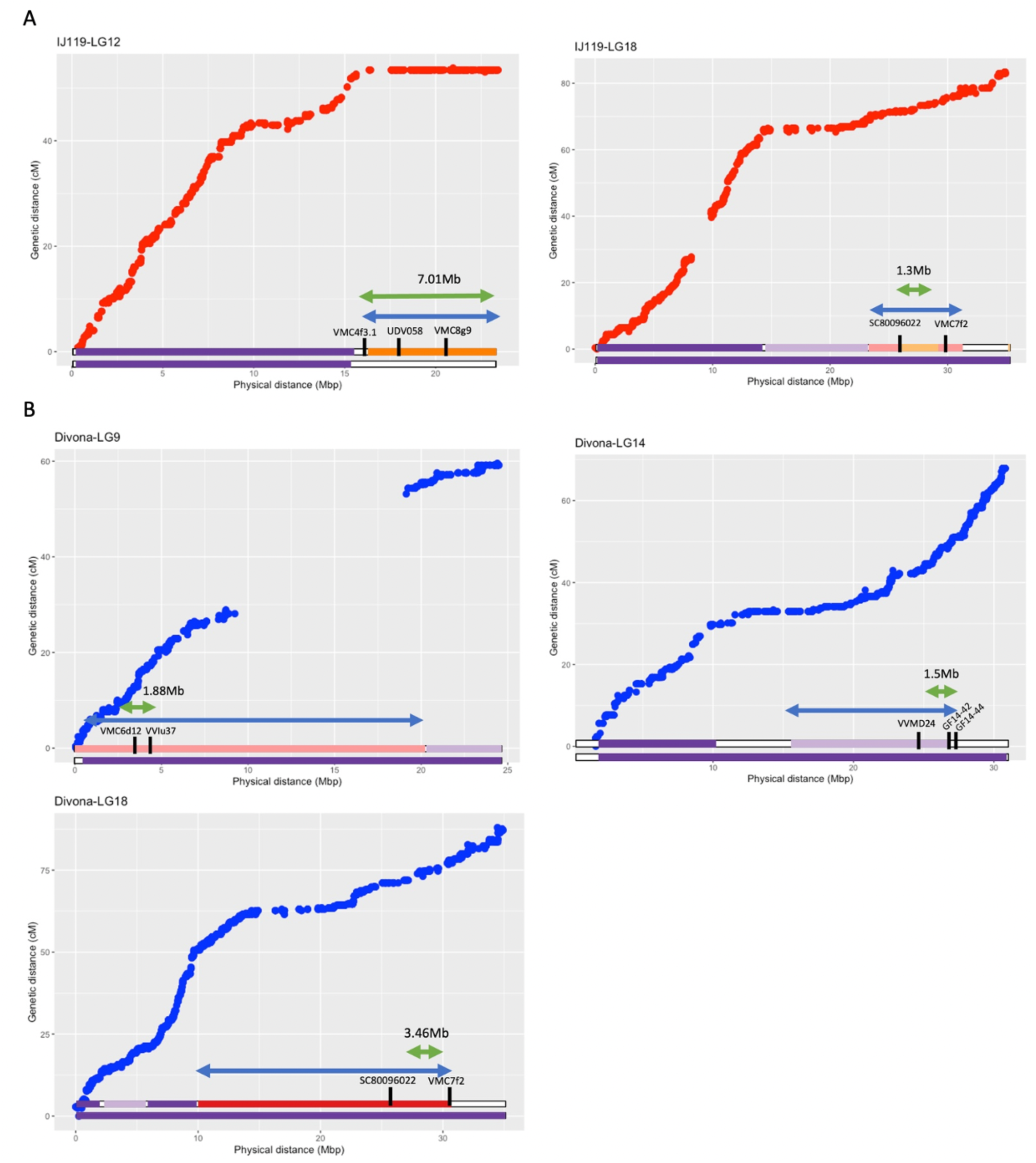
Variation of genetic distance depending on physical distance of markers on parental maps. LG 12 & 18 on IJ119 in red (A). LG9, 14 & 18 on Divona in blue (B). Bars below the graph show the origin of haplotypic blocks. Blue= *V. aestivalis*, Pink= *V. amurensis*, Red = *V. labrusca*, Orange = *V. rotundifolia*, Yellow = *V. coignetiae*, Light Purple= Ripariae (*V. riparia* / *V. rupestris*), Purple = *V. vinifera*, White= Undetermined. Black marks are SSR markers associated with resistance loci. Blue and Green arrow shows the largest/smallest segregating haplotypic block carrying resistance loci.

On LG18 of IJ119, the recombination rate decreases in parallel with the homeologous region. The sizes ofhaplotypic blocks carrying *Rpv3.1* vary between 1.3 and 7.86 Mbp in the 50025 progeny. Surprisingly, chromosome painting of chromosome 18 suggests that this haplotypic block, associated with markers SC80096022 and VMC7f2, derives from the Asian species *V. coignetiae* while the literature on *Rpv3.1* (Di Gaspero et al., 2012) and the known genealogy of IJ119 put forward that the donor species of *Rpv3.1* is American but not Asian.

Divona is an interspecific complex hybrid of *V. vinifera*, *V. amurensis* and American *Vitis* and carries four resistance genes *Rpv10*, *Rpv3.4, Ren3* and *Rgb1* (Figure 4B). The recombination rate on LG9 is constant all along the chromosome. The *Rpv10* is located in a *V. vinifera-V. amurensis* homeologous region. The haplotypic block carrying *Rpv10* ranges in size from 1.88 to 20 Mbp in the progeny.

The QTL *Rgb1* of resistance to black rot is located between 26 and 28 Mbp on LG14. The pedigree of the male parent Divona suggests that *Rgb1* was transmitted from Merzling to Bronner, the grandparent of the population and originates from an American *Vitis*. The chromosome painting of Divona reveals that the haplotypic block carrying *Rgb1* derives from the *Riparae* group (*V. riparia* or *V. rupestris).* The recombination rate in the region carrying *Rgb1* decreases slightly, compared to the rest of the chromosome, resulting in haplotypic blocks of different length in the progeny (min 1.5 Mbp - max 11.26 Mbp).

*Rpv3.3* locus of Divona is located on LG18 between two SSR at 26.8 and 30.5 Mbp. This homeologous region indicates that *Rpv3.3* origin is *V. labrusca*, a native American species. The recombination rate in the homeologous region decrease compared to the homologous region on the same chromosome. The length of the haplotypic block carrying *Rpv3.3* transmitted to the progeny varied between min of 3.46 and max of 20.5 Mbp.

On LG15, the high number of ABxAB type markers results in low SNP density. The lack of relevant data in this region makes chromosome painting not reliable enough, which consequently, was not taken into consideration.

## 4. Discussion

The main objective of our study was to evaluate the effect of the introgression of disease resistance genes from wild *Vitis* on plant performance. Thus, we were able to determine the length of linkage drag around six resistance loci by defining the species of origin of haplotypic blocks in the population of complex interspecific hybrids. The high-quality genetic maps served to decipher the genetic determinism of important agronomical traits in the population and detect several cases of QTL colocalizations between resistance factors and cultural traits.

One of the originalities of this work is the plant material we studied. The population is a breeding population that was obtained as part of INRAE-ResDur breeding program from which several varieties were actually selected for commercial use. This plant material is representative of many breeding materials used in European breeding programs, largely sharing the same disease resistance factors and therefore common genome origins. For this reason, our results can be extended to other populations.

For the selection of new varieties, the breeders aim usually to reduce the percentage of non-vinifera species and the linkage drag around introgressed resistance loci. Linkage drag is usually defined as the “undesirable” effects of genes linked to the introgressed QTL. We compared the variations in recombination rate on LGs carrying resistance loci, taking into consideration the origin of haplotypic blocks. The recombination rate in homeologous regions does not always decreases, it depends on the wild species. As observed, the recombination rate between *Muscadinia* and other *Vitis* species is almost null resulting in a fixed linkage drag with the resistance loci as *Rpv1/Run1* for example. These results are consistent with the conclusion of (Delame et al., 2019). Thus, a high number of progenies is required to break the introgressed block from *V. rotundifolia*. Wild relatives of *V. vinifera* from a closer gene pool like *V. labrusca* and *V. riparia* have less effect on the recombination rate as we observe a decrease in recombination rate but not a total stop as the case of *Muscadinia*. The recombination rate in the homeologous region *V. vinifera* - *V. amurensis* in this population did not change from the homologous regions. These observations mean that the length of linkage drag depends at least partially on the donor species and in most cases can be highly reduced through several backcrossing events like the case of *Rpv3*, *Rgb1,* and *Rpv10*. The size of the haplotypic block carrying *Rpv3.1* varied between 11.5 and 22.6 Mbp in nine resistant varieties while 7.1 Mbp linkage drag was defined around *Rpv1/Run1* loci (Foria et al., 2022).

Our results showed also that the length of linkage drag is not the only factor that should be taken into consideration. Among five resistance loci analyzed in this population, *Rpv1/Run1* from *Muscadinia* had the largest haplotypic block but no QTL was detected in this region for all the studied traits. In tomato, *Tm-2a* resistance gene to ToMV contributed to 64.1 Mb linkage drag (van Rengs et al., 2022) but did not have a high agronomical effect (Rubio et al., 2016).

From a breeding point of view, it’s important to identify colocalizations between QTLs of different traits and resistance genes. This study revealed the colocalization of *Rpv10* with QTL of TSS in berries and pruning wood weight. On the same haplotypic block from *V. amurensis* on LG9, QTL of potassium content in berries and budbreak date are linked to *Rpv10*. The colocalization of QTLs of sugar content in fruits and resistance factor to powdery mildew has been observed also in *Prunus davidiana* (Quilot et al., 2004) a wild relative of the cultivated peach that is used in breeding programs as a source of natural resistance to mildews. The 11.26 Mbp haplotypic block from *V. riparia/V. rupestris* on LG14 carried *Rgb1* along with QTL of number of clusters, berry weight, budbreak date. So, the second colocalization revealed that *Rgb1* is linked to higher number of clusters, earlier genotypes and smaller berries.

Many studies assessed the cost of disease resistance genes on yield, especially in annual plants such as barley or rye (Brown, 2002). Four experiments in barley revealed no significant yield reduction in lines carrying R-genes (Kjaer et al., 1990; Kolster & Stolen, 1987).

The haplotypic block of 11.26 Mbp carrying *Rpv3.3*, carries also two QTLs of yield components and three QTLs of berry composition while the 7 Mbp haplotypic block carrying *Rpv1/Run1* does not have any other QTL for the studied traits. *Rpv3.1* is linked to a QTL of berry weight on the same haplotypic block of 7.86 Mbp. Several studies evaluated the effects of introgressed resistance genes on the performance of cultivated plants such as potato, tomato and peach (Bormann et al., 2004; Brouwer & St Clair, 2004; Chitwood-Brown et al., 2021; Danan et al., 2011; Quilot et al., 2004). Rubio et al. (2016) evaluated the effect of *Tm-2a* resistance gene to Tomato mosaic virus (ToMV), *Sw-5* resistance gene to Tomato spotted wilt virus (TSWV) and *Ty-1* resistance gene to Tomato yellow leaf curl virus (TYLCV) in tomato. ToMV-resistant plant showed higher yield, lower soluble solids and titratable acidity. TSWV-resistant plants had smaller fruit weight. The effects of *Ty-1* were more significant so resistant plants were less productive and had lower acidity and soluble solids. These effects are probably the result of linkage drag around the three resistance genes (Rubio et al., 2016).

Our results revealed the polygenic character of most agronomic traits in grapevine which is consistent with the previously published results (Costantini et al., 2008; Duchêne, Butterlin, et al., 2012; Fechter et al., 2014; Doligez et al., 2013; Bayo-Canha et al., 2019). The most important agronomical traits are controlled by several quantitative loci with a low phenotypic variance explained by each of them. For example, the heat sum requirements at budburst was shown to be controlled by seven loci explaining around 66% of the phenotypic variance. The loci on LG14 colocalized with *Rgb1* and explained only 4.1% of all observed variance. From a breeding point of view, the polygenic nature of cultural traits mitigates the effect of QTL colocalizations with resistance factors. These colocalizations are not of major concern as their effects is relatively low and can be compensated by the presence of other quantitative factors.

The origin of many resistance loci is already known while other are more debatable. In this study, the origin species of the haplotypic blocks carrying *Rpv1/Run1*, *Rpv10* and *Rpv3.3* were consistent with the literature (Di Gaspero et al., 2012; Merdinoglu et al., 2003; Schwander et al., 2012). The haplotypic block of *Rpv3.1* in this study was associated to *V. coignetiae*, an Asian species which is not consistent with genealogy of IJ119. So, the origin of *Rpv3* locus is still under debate. According to the hierarchical clustering within subgenus *Vitis* based on 24 nuclear microsatellites analyzed by Péros et al. (2011), *V. coignetiae* was genetically closer to *Labruscae* group than Asian *Vitis* species. Di Gaspero et al (2012) reported 7 haplotypes of *Rpv3* from five different resistant varieties. The *Rpv3.1* haplotype is known to be introgressed in Chambourcin from Seibel 4614. Foria et al. (2022) found also that the region of *Rpv3.1* matched with Asian species rather than American species and they suggested that *Rpv3.1* is genetically different from all known varieties and originates from a marginal habitat species. *Rgb1* was associated to *Riparae* group (*V. riparia* or *V. rupestris*). The available accessions and sequences that we have are not enough for us to distinguish between these two species. (Rex et al., 2014) detected *Rgb1* in Börner (*V. riparia* Gm183 × *V. cinerea* Arnold) and they assumed that it comes from American *V. cinerea* without having enough arguments to exclude *V. riparia*. Our results strongly suggest that the origin of *Rgb1* is probably *V. riparia*.

Overall, many QTL of agronomical traits were detected in the population and originated from wild species independently from resistance genes. All species, beside *V. rotundifolia*, contributed in the genetic variation of many agronomical traits in the population. We can highlight some traits of interest for the current breeding context that are dominated by the wild species. The American species were at the origin of genetic variability to traits related mainly to yield components (LG11, 14 & 18) and berry composition (LG 14, 17 & 18). The alleles from American *Vitis* leads to smaller berries and higher plant yield. Additionally, two important factors for berry acidity as Potassium and yeast assimilable nitrogen were mainly controlled by American species. Asian species conferred mainly QTLs of phenology on LG7 and on LG9. The Asian alleles induce earlier phenological periods.

Although the crop wild relatives have been used for plant improvement in many perennial and annual species (Hajjar & Hodgkin, 2007; Migicovsky & Myles, 2017), they are usually associated with low quality and negative cultural effects in grapevine. Our results showed that wild species can improve cultural traits and resistance at the same time. For example, the introgression of *Rgb1* for the resistance to black rot is linked to smaller berries and higher cluster number which are two favorable traits for wine production. The favorable and unfavorable traits in the context of grapevine breeding are not simple to define since the ideotype highly depends on the region, the environmental conditions, and the final product. The grapevine wild relatives can indeed be used as sources of natural resistance to biotic stresses but also as sources of adaptation to abiotic stresses arising in the context of climate change.

Through our study, we noticed some particular genetic behaviors in specific LGs. The transition from homologous to a homeologous region on some LGs explains the variation in the recombination rate as on LG12 of IJ119 or LG3 & 10 of Divona (Figure S7 A). But some behaviors were independent of the presence or absence of wild haplotype (Figure S7 B&C). The homeologous structure on LG18 is different between both parents but we see in both cases the same pattern in the recombination rate (Figure 4). A decrease in recombination rate is observed in the second arm of the chromosome which is a region enriched in TIR-NB-LRR R-genes. These observations suggest that the nature of genomic regions can also affect meiotic recombination. Also, the recombination rate in the region between 11 and 19 Mbp on LG19 for both parents dropped significantly and independently from the presence of homologous/homeologous regions (Figure S7 C). This phenomenon has been observed in several genetic maps (Delame et al., 2019; Vervalle et al., 2022).

## Conclusion

To summarize, the chromosome painting analyses conducted on the population of interspecific hybrids confirmed that the risk of linkage drag is higher with *V. rotundifolia* compared to other *Vitis* sp. To our knowledge, no other study has been reported studying the effect of introgression of main disease resistance genes on grapevine performance in the vineyard. The colocalizations of QTL of cultural traits and resistance factors can be overcome due to the polygenic nature of agro-oenological traits. To complete our results and have a complete overview, it is important to conduct metabolomics analyses on the berry juice and wine from this population to detect any specific metabolites from wild species that could be linked to resistance genes and evaluate the effect of wild species on wine quality.

## Author contributions

## Data availability

