## Supplementray information for "Impact of the introgression of resistance loci on agro-œnological traits in grapevine interspecific hybrids"

Supplementary data of "Impact of the introgression of resistance loci on agro-oenological traits in grapevine hybrids"

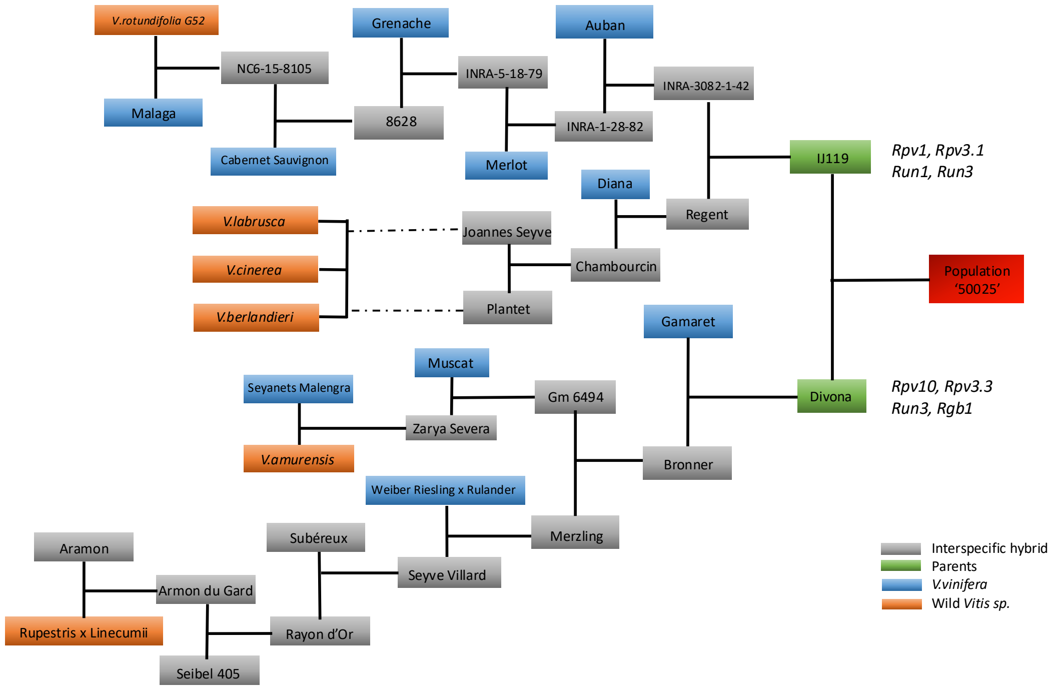


**Figure S1** Part of the genealogy of IJ119 x Divona progeny. IJ119 is the female parent. Divona is the male parent, a white grape variety from Agroscope (Switzerland). *Rpv1, Rpv3.1 Rpv10* and *Rpv3.3* are downy mildew resistance loci. *Run1* and *Ren3* are powdery mildew resistance loci. *Rgb1* is a black rot resistance locus.

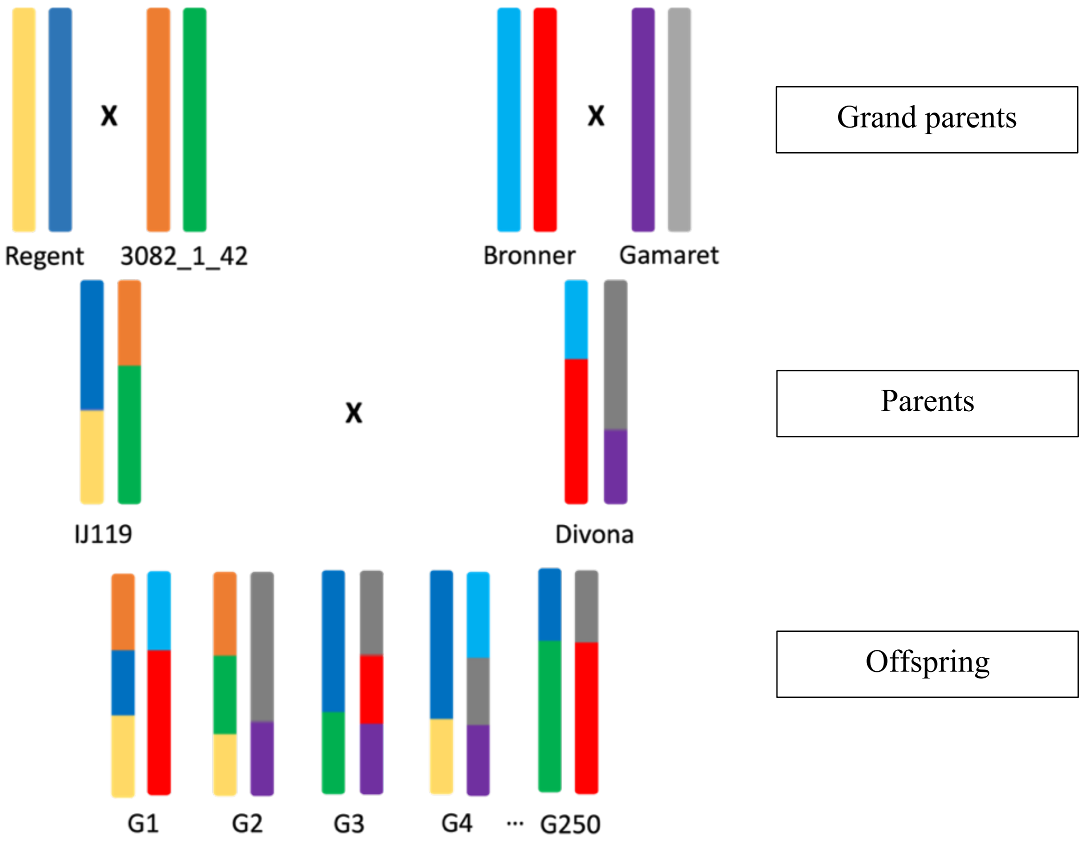


**Figure S2** Visual illustration of the principles of Chromosome painting analysis

**Table S1** Heritabilities of phenotypic traits

| | Variable | Vitis ontology | Abbreviation | Heritabilities | | |
|---|---|---|---|---|---|---|
| | | | | 2019 | 2020 | 2021 |
| *Phenology* | Heat sum base 2 from 15 February to budburst | CO_356:1000088 | HS_B | 0.50 | 0.77 | 0.79 |
| | Heat sum base 10 from budburst to flowering | CO_356:1000090 | HS_BF | 0.55 | 0.58 | 0.57 |
| | Heat sum base 6 from flowering to véraison | CO_356:1000089 | HS_FV | 0.81 | 0.96 | 0.98 |
| *Vigor* | Chlorophyll content | CO_356:1000332 | SPAD | NA | 0.68 | 0.93 |
| | Exposed leaf area | CO_356:1000080 | ELA | 0.84 | 0.44 | 0.16 |
| | Pruning wood weight | CO_356:1000190 | PWW | 0.87 | 0.34 | 0.9 |
| | Apparent canopy volume – LiDAR | CO_356:1000540 | ACV | NA | 0.79 | 0.66 |
| | Apparent wood volume – LiDAR | CO_356:1000541 | AWV | NA | 0.83 | 0.81 |
| | Potassium content in petioles | CO_356:1000333 | Pet_K | NA | 0.76 | - |
| | Magnesium content in petioles | CO_356:1000334 | Pet_Mg | NA | 0.44 | - |
| | Bud break ratio | - | BBr | 0.72 | - | - |
| | Secondary shoots ratio | - | SSr | 0.88 | 0.8 | 0.67 |
| | Early leaf yellowing | - | LeafY | NA | 1 | 1 |
| *Yield components* | Number of inflorescences per primary shoot | CO_356:1000267 | Fert_Prim | 0.83 | 0.96 | 0.67 |
| | Number of inflorescences per shoot | CO_356:1000266 | Fert_Tot | 0.86 | 0.78 | 0.75 |
| | Number of bunches per plant | CO_356:1000158 | Clust_Nb | 0.4 | 0.4 | 0.51 |
| | Total bunches weight per plant | CO_356:1000247 | ClustW | 0.95 | 0.62 | 0.76 |
| | Green berry mean weight | CO_356:1000336 | GBW | NA | 0.92 | 0.89 |
| | Berry mean weight | CO_356:1000215 | BW | 0.54 | 0.68 | 0.9 |
| | Fruit set quality | | FruitSet | NA | 0.51 | 0.99 |
| | Bunch compactness | CO_356:1000037 | ClustComp | NA | NA | 0.53 |
| *Berry composition* | Green berries pH | CO_356:1000329 | GB_pH | NA | 0.83 | 0.97 |
| | Berries pH | CO_356:1000184 | B_pH | 0.83 | 0.93 | 0.66 |
| | Berries titratable acidity | CO_356:1000231 | B_TA | 0.47 | 0.73 | 0.28 |
| | Berries total soluble solids | CO_356:1000239 | B_IR | 0.26 | 0.73 | 0.97 |
| | Green berries malic acid content | CO_356:1000325 | GB_Mal | NA | 0.39 | 0.77 |

| | | | | | |
|---|---|---|---|---|---|
| Berries malic acid content | CO_356:1000298 | B_Mal | 0.56 | 0.15 | 0.45 |
| Green berries tartaric acid content | CO_356:1000326 | GB_Tar | NA | 0.94 | 0.84 |
| Berries tartaric acid content | CO_356:1000300 | B_Tar | 0.42 | 0.22 | 0.86 |
| Green berries potassium content | CO_356:1000330 | GB_K | NA | 0.29 | 0.9 |
| Berries potassium content | CO_356:1000187 | B_K | 0.66 | 0.72 | 0.75 |
| Berries ammonium content | CO_356:1000352 | B_NH4 | 0.01 | 0.63 | NA |
| Green berries yeast assimilable nitrogen content | CO_356:1000539 | GB_YAN | NA | 0.61 | 0.81 |
| Green berries ammonium content | CO_356:1000327 | GB_NH4 | NA | 0.54 | 0.82 |

**Table S2** BLUP of three seasons of the progeny, parents and Chardonnay variety.

| | | | Progeny | | | Divona | IJ119 | Chardonnay | | | |
|---|---|---|---|---|---|---|---|---|---|---|---|
| | Variable | Unit | Min | Mean | Max | Mean | Mean | Mean | Var 2019 | Var 2020 | Var 2021 |
| ***Phenology*** | HS_B | Degree. days | 655.9 | 752.7 | 956.4 | 716.2 | 815.1 | 743.9 | 1543.6 | 842.7 | 607.2 |
| | HS_BF | Degree. days | 521.2 | 573.8 | 627.9 | 579.3 | 561.6 | 582 | 183.2 | 347.1 | 174.5 |
| | HS_FV | Degree. days | 924.7 | 1159.5 | 1430.1 | 992 | 1319.3 | 1158.5 | 1572.3 | 664.8 | 352.6 |
| ***Vigor*** | SPAD | Dimensionless | 269.2 | 318.5 | 364.5 | 291.2 | 296.3 | 313.9 | NA | 268.3 | 51.3 |
| | ELA | $m^2/m^2$ | 1.3 | 2 | 2.5 | 2.1 | 1.8 | 1.9 | 0.02 | 0.02 | 0.04 |
| | PWW | kg/plant | 0.2 | 0.7 | 1.3 | 0.4 | 0.4 | 0.6 | 0.004 | 0.029 | 0.004 |
| | ACV | l/plant | 217.3 | 335.5 | 874.1 | 317.8 | 291.2 | 307.9 | NA | 393.3 | 455.9 |
| | AWV | /plant | 1.85 | 4.28 | 7 | 3.4 | 2.6 | 3.3 | NA | 0.122 | 0.113 |
| | Pet_K | g/100g | 0.9 | 2 | 3.9 | 2.4 | 1.5 | 1.9 | NA | 0.078 | 0.73 |
| | Pet_Mg | g/100g | 0.3 | 0.5 | 0.8 | 0.3 | 0.5 | 0.5 | NA | 0.01 | 0.03 |
| | BBr | % | 44.3 | 81.6 | 98 | 84.4 | 80.6 | 80.5 | 13.1 | 45 | 48.2 |
| | SSr | | 0.04 | 0.12 | 0.39 | 0.34 | 0.04 | 0.08 | 0.002 | 0.005 | 0.001 |
| | LeafY | 1 to 9 | 1 | 2.8 | 7 | 3 | 8 | 1 | NA | 0 | 0 |
| ***Yield components*** | Fert_Prim | per shoot | 1.5 | 2.1 | 2.7 | 2.6 | 2 | 1.8 | 0.01 | 0.001 | 0.001 |
| | Fert_Tot | per shoot | 1.8 | 2.4 | 3.4 | 3.3 | 2.2 | 2 | 0.02 | 0.01 | 0.001 |
| | Clust_Nb | per plant | 21 | 32.4 | 44.8 | 37.2 | 26.6 | 30.6 | 31.1 | 6.6 | 8.3 |
| | ClustW | kg/plant | 1.9 | 4.3 | 6.6 | 4.89 | 3.7 | 3.4 | 0.06 | 0.19 | 0.09 |
| | GBW | g | 0.4 | 0.7 | 1 | 0.7 | 0.7 | 0.8 | NA | $8.10^{-4}$ | 0.002 |
| | BW | g | 0.8 | 1.2 | 2 | 1.1 | 1.1 | 1.3 | 0.02 | 0.01 | 0.008 |
| | FruitSet | 1 to 4 | 0.4 | 1.9 | 2.5 | 2.1 | 2 | 2.1 | NA | 0.03 | 0 |
| | ClustComp | 1 to 9 | 0.6 | 4.9 | 7.9 | 3.8 | 5 | 5.7 | NA | NA | 0.46 |

| | | | | | | | | | | |
|---|---|---|---|---|---|---|---|---|---|---|
| | GB_pH | Dimensionless | 2.3 | 2.4 | 2.5 | 2.5 | 2.4 | 2.4 | NA | $6x10^{-4}$ | $8x10^{-5}$ |
| | B_pH | Dimensionless | 2.8 | 2.9 | 3 | 2.9 | 2.9 | 2.9 | $7x10^{-4}$ | $3x10^{-4}$ | 0.002 |
| | B_TA | meq/l | 120.7 | 145.2 | 172.9 | 138.5 | 151.1 | 155.1 | 0.4 | 0.1 | 0.3 |
| | B_IR | °Brix | 15.4 | 17.3 | 22.5 | 16.7 | 16.9 | 17.7 | 1.02 | 0.25 | 0.04 |
| | GB_Mal | mmol/l | 168.5 | 195.1 | 222.2 | 169.2 | 214.2 | 207.7 | NA | 76.05 | 14.44 |
| | B_Mal | mmol/l | 14.7 | 29.4 | 41.7 | 22 | 37.3 | 38.4 | 4.23 | 12.77 | 6.32 |
| | GB_Tar | mmol/l | 67.5 | 90.8 | 108.1 | 85.8 | 85.3 | 76.8 | NA | 2.33 | 3.25 |
| | B_Tar | mmol/l | 40.9 | 51.6 | 58.8 | 53 | 50.7 | 49.3 | 3.88 | 8.23 | 0.54 |
| | GB_K | mmol/l | 22.6 | 27.3 | 33.7 | 29.4 | 26.2 | 26.6 | NA | 3.67 | 0.17 |
| | B_K | mmol/l | 22.9 | 29.2 | 33.9 | 29.5 | 32.1 | 29.4 | 0.86 | 1.29 | 0.61 |
| | B_NH4 | mg/l | 0.8 | 1.4 | 4.2 | 1.4 | 1 | 1.5 | 0.15 | 0.18 | NA |
| | GB_YAN | mg/l | 184.1 | 259.6 | 407.3 | 140.6 | 134 | 272.9 | NA | 363.3 | 196 |
| | GB_NH4 | mg/l | 54.8 | 125.4 | 269.8 | 125.2 | 94 | 154.5 | NA | 153.3 | 171.6 |

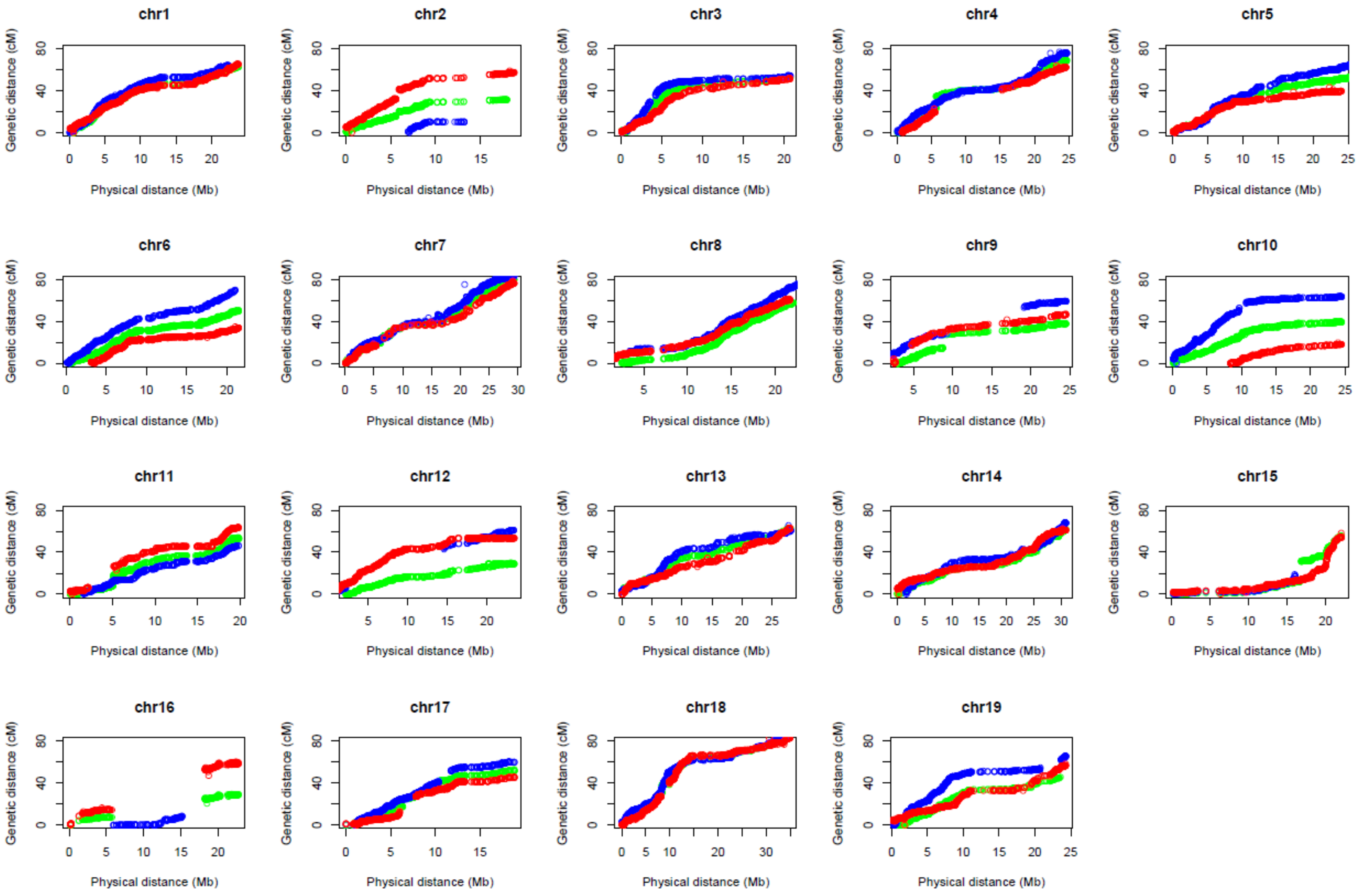


**Figure S3** Comparison between genetic and physical marker order in the linkage groups of three constructed maps with Lep-Map3. The x-axis indicates the position of the markers on the reference genome. Each dot indicates a marker, and its color indicates the map: red for the IJ119 map, blue for the Divona map, and green for the consensus map

**Table S3** Characteristics of QTL detected on the consensus genetic map of the population in study

| | Variable | *LG* | *Position (cM)* | *LOD p=0.05* | *LOD Max* | *% Var* | *CI- Prob=0.95* | |
|---|---|---|---|---|---|---|---|---|
| | | | | | | | *Genetic Pos (cM)* | *PN40024 (Mb)* |
| ***Phenology*** | *HS_B* | 1 | 57.43 | 4.41 | 15.32 | 14.73 | 56.4-60.8 | 21.08-22.9 |
| | | 5 | 49.60 | 4.41 | 15.07 | 14.44 | 48.2-50 | 20.4-22.9 |
| | | 6 | 37.15 | 4.41 | 17.57 | 17.36 | 30.9-37.2 | 9.04-15.58 |
| | | 7 | 17.67 | 4.41 | 7.57 | 6.65 | 3.6-18.3 | 0.57-4.7 |
| | | 9 | 38.40 | 4.41 | 4.85 | 4.13 | 0.4-39.4 | 3.09-23.5 |
| | | 14 | 44.38 | 4.41 | 5.92 | 5.10 | 35.5-50.6 | 22.4-27.3 |
| | | 17 | 4.02 | 4.41 | 4.72 | 4.01 | 0-11 | 0.04-4.46 |
| | *HS_BF* | 1 | 46.19 | 4.3 | 4.657 | 5.29 | 43.1-53 | 10.88-19 |
| | | 6 | 39.36 | 4.3 | 14.88 | 19.03 | 26.51-43.6 | 7.72-19 |
| | | 7 | 13.46 | 4.3 | 6.375 | 7.38 | 9.64-19.1 | 1.78-5.16 |
| | | 8 | 24.50 | 4.3 | 5.932 | 6.84 | 10.84-31.9 | 10.67-15.59 |
| | | 14 | 62.26 | 4.3 | 5.33 | 6.1 | 43.58-62.26 | 25.4-30.8 |
| | *HS_FV* | 1 | 38.56 | 4.50 | 6.24 | 5.23 | 29.52-41.37 | 6.87-10.21 |
| | | 5 | 49.80 | 4.50 | 7.65 | 6.51 | 47.59-52.61 | 19.93-25.47 |
| | | 8 | 18.63 | 4.50 | 6.70 | 5.65 | 10.84-28.11 | 10.67-14.59 |
| | | 14 | 50.81 | 4.50 | 6.94 | 5.86 | 36.15-55.83 | 22.55-28.75 |
| | | 15 | 32.24 | 4.50 | 5.86 | 4.89 | 13.66-53.33 | 16.02-21.5 |
| | | 16 | 27.04 | 4.50 | 29.63 | 32.80 | 8.04-28.85 | 5.39-19.98 |
| ***Vigor*** | *ELA* | 1 | 19.88 | 4.513 | 4.76 | 9.06 | 11.8-41.2 | 3.09-10.1 |
| | | 10 | 39.56 | 4.513 | 5.62 | 10.8 | 38.4-39.6 | 22.7-23.7 |
| | *SPAD* | 7 | 73.71 | 4.45 | 7.60 | 14.26 | 71.5-77.72 | 25.8-28.1 |
| | | 13 | 23.70 | 4.45 | 5.15 | 9.40 | 5.42-41.57 | 0.6-16 |
| | *PWW* | 1 | 19.48 | 4.57 | 4.94 | 8.5 | 7.43-25.7 | 2.03-5.3 |
| | | 9 | 1.21 | 4.57 | 5.55 | 9.6 | 0.4-7.03 | 2.93-4.98 |
| | | 18 | 59.08 | 4.57 | 6.03 | 10.45 | 52-70.32 | 11.3-25.5 |
| | *ALV* | 1 | 38.96 | 4.35 | 9.17 | 15.94 | 7.43-41.57 | 2-10.2 |
| | | 3 | 7.43 | 4.35 | 4.64 | 7.62 | 0-16.27 | 0.19-2.9 |
| | | 10 | 38.56 | 4.35 | 8.97 | 15.56 | 36.95-39.56 | 17.8-23.7 |
| | *AWV* | 1 | 32.53 | 4.33 | 6.03 | 10.37 | 11.85-36.55 | 3.09-8.4 |
| | | 9 | 2.21 | 4.33 | 4.32 | 7.28 | 0.8-10.04 | 3.13-6.05 |
| | | 10 | 38.56 | 4.33 | 8.01 | 14.08 | 34.74-39.56 | 14.7-23.58 |
| | *Pet_K* | 6 | 20.28 | 4.46 | 5.37 | 9.6 | 12.45-31.73 | 4.7-10.8 |
| | | 7 | 68.49 | 4.46 | 6.68 | 12.1 | 63.67-73.31 | 23.1-26.5 |
| | *Pet_Mg* | 18 | 17.2 | 4.44 | 4.66 | 9.8 | 6.23-26.31 | 0.93-7.5 |
| | *Pet_K/Mg* | 6 | 28.9 | 4.38 | 5.98 | 11.25 | 25.3-32.13 | 7.35-10.79 |
| | | 7 | 68.8 | 4.38 | 4.1 | 7.53 | 65.67-77.52 | 23.6-28.08 |
| | *LeafY* | 10 | 37.35 | 4.46 | 7.09 | 10.9 | 33.54-38.36 | 12.77-22.77 |
| | | 16 | 27.04 | 4.46 | 4.60 | 7.9 | 4.22-32.46 | 12.93-21.82 |
| | | 18 | 58.87 | 4.46 | 5.64 | 8.7 | 24.70-66.31 | 7.28-22.95 |
| | *SSr* | 1 | 24.90 | 4.4 | 13.89 | 20.55 | 11.8-32.3 | 3.08-7.4 |
| | | 5 | 43.18 | 4.4 | 9.60 | 13.51 | 35.1-46.1 | 11.2-19.4 |
| | | 6 | 22.09 | 4.4 | 6.07 | 8.19 | 12.8-29.32 | 4.7-8.4 |
| | | 18 | 9.24 | 4.4 | 4.92 | 6.56 | 0-18.4 | 4.4-5.9 |
| | *BBr* | 3 | 16.47 | 4.39 | 6.02 | 10.53 | 14.6-19.6 | 2.7-3.4 |
| | | 17 | 10.44 | 4.39 | 4.93 | 8.5 | 0-25.75 | 0.07-6.8 |
| ***Yield co*** | *Fert_prim* | 8 | 36.95 | 4.42 | 5.28 | 9.15 | 35.1-53.8 | 16.5-20.7 |
| | | 14 | 30.12 | 4.42 | 9.26 | 16.8 | 25.7-33.5 | 12-21.4 |

| | | | | | | | | |
|---|---|---|---|---|---|---|---|---|
| | *Fert_tot* | 14 | 30.12 | 4.49 | 7.95 | 16.4 | 25.7-35.5 | 12-22.4 |
| | *Clust_Nb* | 1 | 39.96 | 4.37 | 6.17 | 10.99 | 37.5-48.4 | 8.6-17.7 |
| | | 14 | 46.59 | 4.37 | 8.29 | 15.12 | 32.7-56.6 | 21.5-29.2 |
| | *ClustW* | 1 | 61.85 | 4.32 | 5.88 | 9.18 | 27.1-62.6 | 5.7-23.6 |
| | | 17 | 29.97 | 4.32 | 6.70 | 10.59 | 17.6-42 | 6.2-10.5 |
| | | 18 | 58.07 | 4.32 | 6.95 | 11.03 | 54.2-59.8 | 11.8-13.1 |
| | *GBW* | 1 | 42.57 | 4.4 | 8.08 | 8.57 | 39-51.2 | 9.3-18.8 |
| | | 8 | 24.50 | 4.4 | 8.95 | 9.6 | 13.4-27.5 | 11.8-14.4 |
| | | 11 | 4.42 | 4.4 | 7.88 | 8.34 | 0-24.3 | 2.8-7.7 |
| | | 14 | 44.18 | 4.4 | 8.31 | 8.85 | 36.5-54 | 22.5-28.3 |
| | | 17 | 25.35 | 4.4 | 12.31 | 13.73 | 17.7-28.8 | 6.2-8 |
| | *BW* | 1 | 56.43 | 4.46 | 5.97 | 5.63 | 51.2-60.4 | 18.8-22.3 |
| | | 8 | 19.08 | 4.46 | 10.45 | 10.4 | 13.2-20.2 | 11.8-13.4 |
| | | 11 | 23.17 | 4.46 | 12.69 | 12.96 | 20.3-25.3 | 6.4-7.9 |
| | | 14 | 40.17 | 4.46 | 10.96 | 10.97 | 35.7-52.2 | 22.5-27.9 |
| | | 17 | 15.66 | 4.46 | 4.70 | 4.37 | 14.4-32.3 | 5.5-9 |
| | | 18 | 62.89 | 4.46 | 8.53 | 8.29 | 59.8-69 | 13.1-23.8 |
| | *FruitSet* | 13 | 10.64 | 4.38 | 5.5 | 11.7 | 4.21-30 | 0.5-8 |
| | *ClustComp* | 11 | 16.14 | 4.3 | 4.75 | 8.94 | 7.6-31 | 4.9-10 |
| | | 12 | 16.87 | 4.3 | 4.3 | 8.1 | 10.5-22.8 | 6.6-16.3 |
| **Berry composition** | *B_IR* | 1 | 25.70 | 4.48 | 4.84 | 6.40 | 6.8-62.6 | 1.8-23.6 |
| | | 9 | 1.41 | 4.48 | 8.23 | 11.3 | 1.2-3.6 | 3.3-4.1 |
| | | 16 | 28.45 | 4.48 | 8.74 | 12.0 | 8.04-29.05 | 5.39-20 |
| | | 17 | 4.02 | 4.48 | 4.96 | 6.50 | 0-13.25 | 0.07-5.2 |
| | | 18 | 58.07 | 4.48 | 5.10 | 6.70 | 52.8-63.1 | 11.4-14.6 |
| | *GB_pH* | 5 | 13.45 | 4.37 | 6.44 | 9.46 | 6.23-47.19 | 2.4-19.69 |
| | | 13 | 50.81 | 4.37 | 4.95 | 7.15 | 36.95-62.06 | 13.09-27.96 |
| | | 14 | 34.74 | 4.37 | 7.76 | 11.58 | 14.86-45.19 | 6.4-25.85 |
| | | 18 | 53.65 | 4.37 | 6.84 | 10.09 | 50.84-58.07 | 11.24-12.49 |
| | *B_pH* | 1 | 41.97 | 4.39 | 4.97 | 11.15 | 5-44.38 | 0.47-11.53 |
| | | 6 | 1.41 | 4.39 | 4.27 | 7.6 | 0.6-8.43 | 0.57-3.46 |
| | *B_TA* | 6 | 1.21 | 4.45 | 4.58 | 7.24 | 0-11.05 | 0.53-4.2 |
| | | 14 | 33.34 | 4.45 | 6.45 | 10.43 | 12.85-42.78 | 4.74-25.47 |
| | | 17 | 4.42 | 4.45 | 7.61 | 12.46 | 1.41-10.4 | 1.24-4.29 |
| | *GB_Mal* | 5 | 15.00 | 4.41 | 5.28 | 8.6 | 3.82-46.19 | 0.78-19.30 |
| | | 13 | 7.23 | 4.41 | 8.93 | 15.18 | 5.42-10.24 | 0.68-2.78 |
| | *B_Mal* | 6 | 21.09 | 4.37 | 4.37 | 9.17 | 1.21-43.38 | 0.87-18.98 |
| | *B_Tar* | 17 | 2.21 | 4.32 | 4.34 | 9.21 | 0-30.28 | 0.007-17.2 |
| | *B_Tar/Mal* | 6 | 22.29 | 4.21 | 4.67 | 10 | 8.03-38.16 | 3.29-16.85 |
| | *GB_K* | 6 | 23.50 | 4.4 | 4.49 | 8.75 | 15.86-40.77 | 5.31-18.04 |
| | | 11 | 25.38 | 4.4 | 5.13 | 10.07 | 3.61-27.99 | 2.6-8.5 |
| | *B_K* | 9 | 0.40 | 4.44 | 6.14 | 11.70 | 0.4-2.2 | 2.93-3.65 |
| | | 17 | 2.61 | 4.44 | 5.09 | 9.59 | 0-3.41 | 0.007-17.2 |
| | *GB_YAN* | 17 | 11.45 | 4.47 | 7.00 | 10.08 | 7.43-15.46 | 3.1-5.79 |
| | *GB_NH4* | 3 | 37.36 | 4.37 | 6.62 | 11.11 | 16.87-45.39 | 3.03-9.44 |
| | | 14 | 33.94 | 4.37 | 5.33 | 8.83 | 28.31-36.14 | 19.23-22.55 |
| | | 17 | 10.84 | 4.37 | 6.78 | 11.4 | 7.43-15.46 | 3.1-5.79 |

**Table S4** Characteristics of QTL of K/Mg detected on IJ119 genetic map.

| *Variable* | *LG* | *Lod à p=0.05* | *Lod max* | *Position (cM)* | *IC à P=0.95* cM | Mbp | *% Var* |
|---|---|---|---|---|---|---|---|
| ***K/Mg_2Y*** | 6 | 2.84 | 6.11 | 21.69 | 4.82-26.11 | 4.84-15.43 | 11.33 |
| | 10 | 2.84 | 3.98 | 6.02 | 2.81-12.04 | 9.92-14.71 | 7.20 |
| | 14 | 2.84 | 3.29 | 25.31 | 9.65-37.36 | 0.98-22.33 | 5.91 |

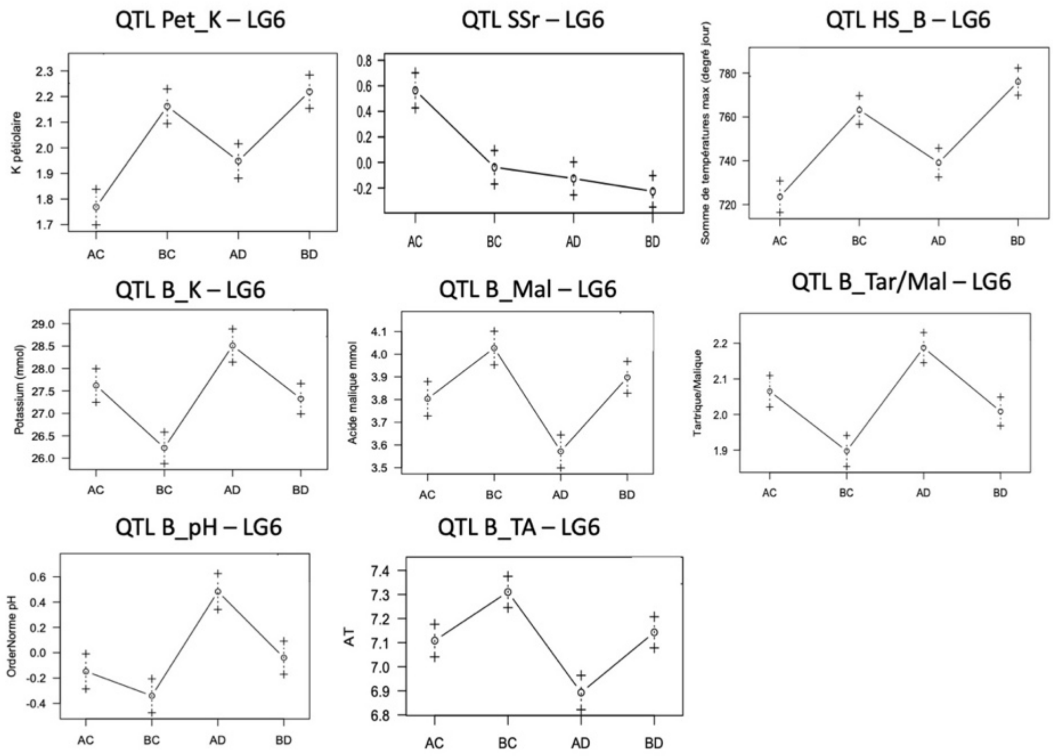


**Figure S4** Plots presenting allele effects of agronomical QTLs on LG6. AB (IJ119) x CD (Divona).

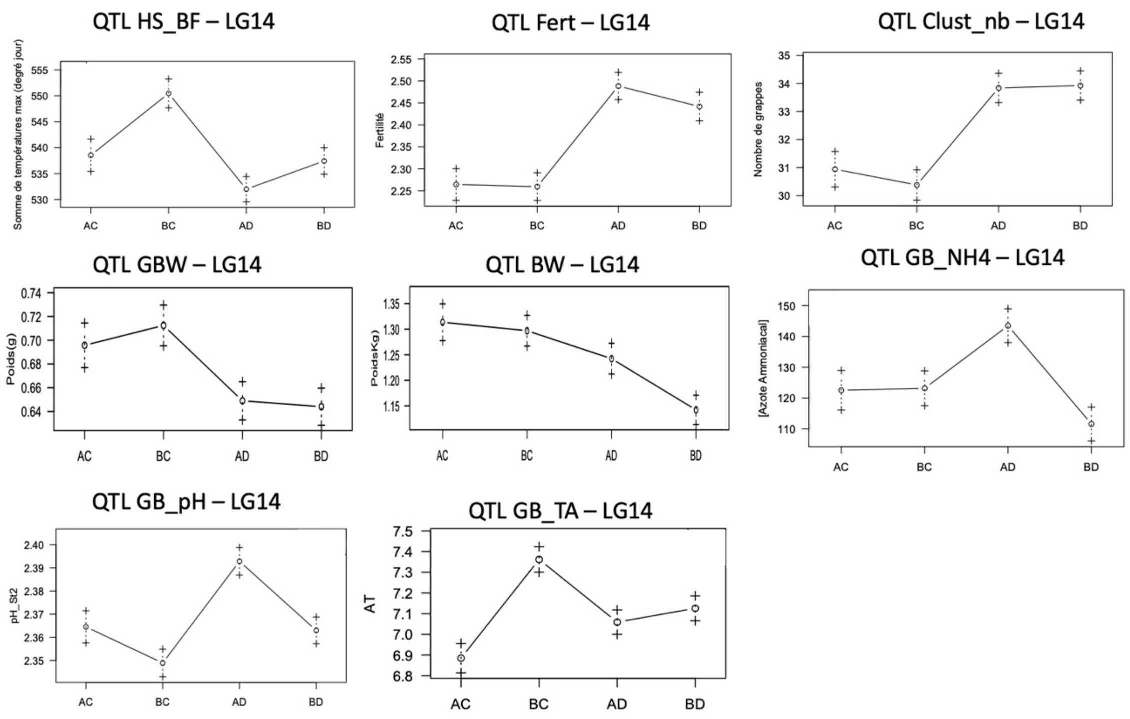


**Figure S5** Plots presenting allele effects of agronomical QTLs on LG14. AB (IJ119) x CD (Divona).

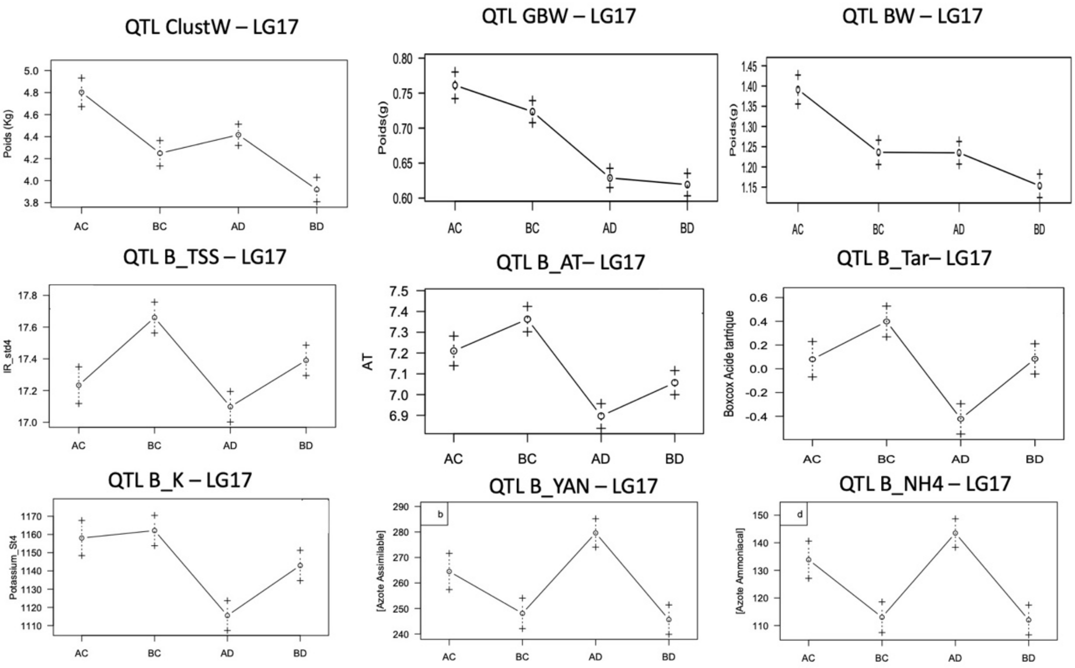


**Figure S6** Plots presenting allele effects of agronomical QTLs on LG17. AB (IJ119) x CD (Divona)

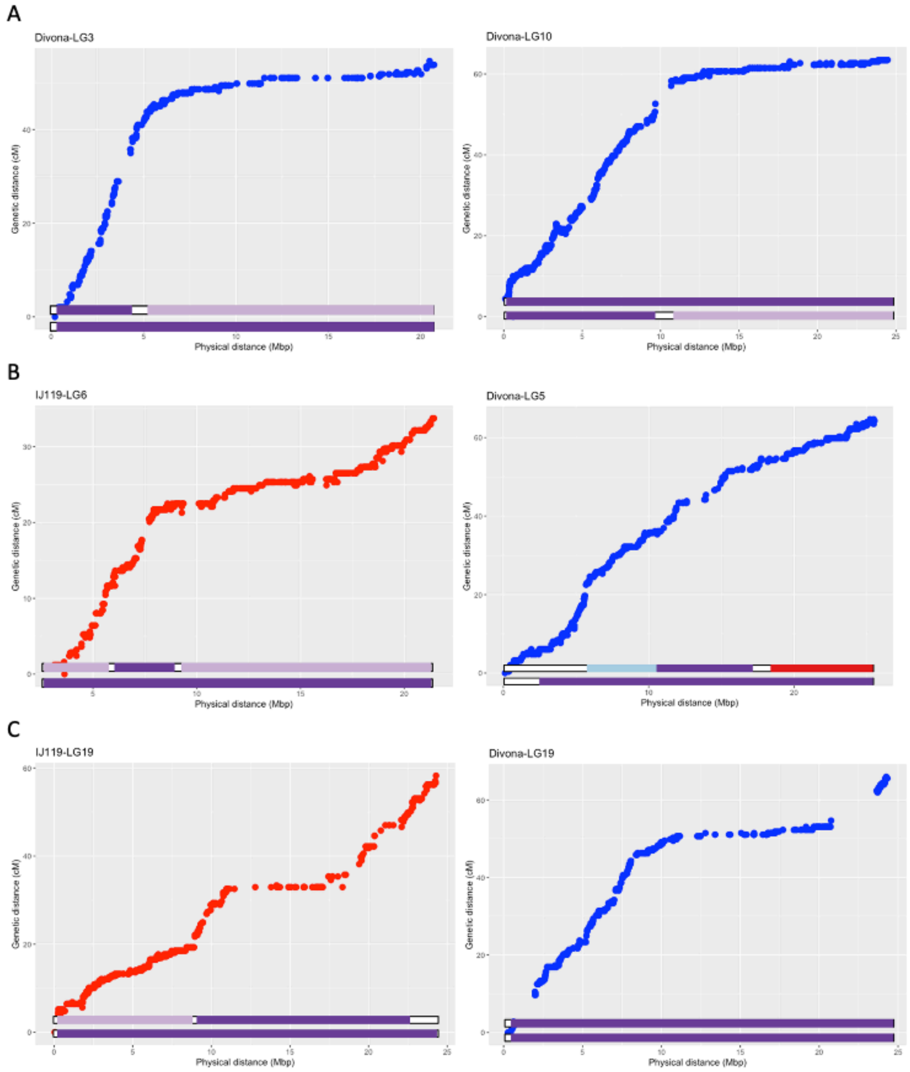


**Figure S7** Variation of genetic distance depending on physical distance of markers on parental maps. LG 6 & 19 on IJ119 in red. LG3, 5, 10 & 19 on Divona in blue. Bars below the graph show the origin of haplotypic blocks. Blue= *V. aestivalis*, Pink= *V. amurensis*, Red= *V. labrusca*, Orange = *V. rotundifolia*, Yellow = *V. coignetiae*, light Purple= Ripariae (*V.riparia / V.rupestris*), Purple = *V. vinifera*, white= Undetermined
